# Affinity Maturation Enables Dual Targeting of Cancer-Associated MUC1-Tn/STn and Drives Potent Tumor Control

**DOI:** 10.64898/2026.10.22.761102

**Authors:** Jinfeng Ye, Xueyong Zhu, Huinan Lu, Jessica Coppola, Ding Zhang, Yinping Tian, Zhuo Yang, Shereen Chung, Leigh-Anne Cui, Jinghao Zhi, Liuqing Wen, Jianzhong Jeff Xi, Ian A. Wilson, Joseph Jardine, Peng Wu

## Abstract

A major challenge in solid tumor immunotherapy is identifying tumor-restricted antigens while overcoming intratumoral antigen heterogeneity. The truncated O-glycans Tn (GalNAcα-O-Ser/Thr) and STn (Neu5Acα2-6GalNAcα-O-Ser/Thr) are aberrantly displayed on the transmembrane glycoprotein MUC1 across a broad spectrum of epithelial malignancies, making MUC1-Tn/STn an attractive shared tumor-associated antigen class. However, heterogeneous intratumoral Tn/STn expression renders single-glycoform-specific antibodies vulnerable to antigen-driven escape. To address this challenge, we used structure-guided directed evolution and yeast surface display to affinity-mature the anti-MUC1-Tn antibody 5E5, generating variants with high-affinity dual recognition of MUC1-Tn and MUC1-STn. Co-crystal structures revealed that heavy-chain CDR loops cooperatively engage the GalNAc moiety primarily through hydrogen-bonding networks, in which HCDR3 residue 98 fine-tunes glycoform preference: Tyr98 favors MUC1-Tn recognition, whereas His98 enhances MUC1-STn engagement, while both support high-affinity recognition of both glycoforms. When reformatted as bispecific T-cell engagers, the affinity-matured variants mediated potent and durable cytolytic activity against cancer cells with heterogeneous glycan profiles, suppressed mixed Tn/STn tumors in vivo where the parental antibody failed, and showed enhanced activity in patient-derived tumor-like clusters. Together, these findings establish functional glycoform coverage as a design principle for antibody engineering, whereby a single paratope is optimized to recognize a defined spectrum of tumor-associated glycoforms while preserving antigen specificity, providing a strategy to overcome glycan heterogeneity in solid tumor immunotherapy.

**Highlights:**

- Directed evolution of antibody 5E5 generates variants with high-affinity dual recognition of MUC1-Tn and MUC1-STn.
- HCDR3 98 acts as a fine-tuning mechanism for glycoform preference, Tyr98 preferentially favors the recognition of MUC1-Tn, His98 shifts this preference toward MUC1-STn, whilst both residues support high-affinity binding to both glycoforms.
- A broadly transferable W27d substitution enhances glycopeptide recognition across structurally distinct anti-MUC1-Tn/STn antibody scaffolds.
- Engineered bispecific T-cell engagers overcome intratumoral glycan heterogeneity, suppressing tumor growth where the parental antibody fails.

## Introduction

Truncated O-glycans are among the most tumor-restricted antigens identified in human epithelial malignancies. The Tn antigen (GalNAcα1-O-Ser/Thr) and its sialylated derivative STn (Neu5Acα2-6GalNAcα1-O-Ser/Thr) are virtually absent from normal adult epithelia but aberrantly overexpressed across breast, pancreatic, ovarian, lung, and colorectal carcinomas, establishing them as well-validated tumor-associated carbohydrate antigens (TACAs)(1–5). Their accumulation is mechanistically rooted in their glycan biosynthetic defects: loss-of-function mutation or epigenetic silencing of *C1GALT1* and its chaperone COSMC impairs the elongation of GalNAc residues beyond the initial sugar, while concurrent upregulation of *ST6GALNAC1* drives direct α2,6-sialylation of GalNAc, prematurely terminating O-glycan chain extension(1,6–9). These mechanisms are not mutually exclusive, individual tumor cells within the same lesion characteristically co-express both glycoforms at variable and shifting ratios, a pattern of intratumoral glycoform heterogeneity that renders single-glycoform-specific therapeutic strategies inherently susceptible to antigen-driven immune escape(1,2,8,10).

MUC1 is a major oncogenic carrier of the tumor-associated Tn and STn glycoforms in epithelial malignancies. Its extracellular variable numbers of tandem repeats (VNTRs) domain contains 20-residue repeats harboring three antibody-accessible epitopes: GVTS, PDTR, and GSTA(11–14). In normal glandular epithelia, MUC1 is largely confined to the apical surface and densely decorated with elongated O-glycans that contribute to epithelial barrier function(11,15,16). Malignant transformation disrupts this organization: MUC1 becomes overexpressed and redistributed across the tumor cell surface, while defective O-glycan biosynthesis promotes accumulation of Tn and STn. In parallel, overexpressed MUC1 engages oncogenic signaling pathways, including NF-κB and EGFR networks, to promote tumor progression and therapeutic resistance. (1,14,17–19). MUC1-Tn/STn also contributes to immune evasion through glycan–lectin interactions, including engagement of MGL/CD301 and inhibitory Siglec pathways. (20–24). Together, these oncogenic and immunosuppressive properties help explain the association of MUC1-Tn/STn with poor clinical outcomes and make aberrantly glycosylated MUC1 an attractive target for cancer immunotherapy. (3–5).

The tumor-restricted display of Tn/STn on MUC1 has motivated therapeutic development across multiple modalities over three decades. MUC1-based glycopeptide and carbohydrate-conjugate vaccines can induce antigen-specific humoral responses but have not produced consistent improvements in patient survival, reflecting the intrinsic challenges of carbohydrate antigens, including weak immunogenicity, tolerance to self-associated mucin glycans, and difficulty eliciting durable cytotoxic T cell responses (25–30). More recently, Daiichi Sankyo developed DS-3939a, an antibody-drug conjugates (ADCs) targeting the PDTR epitope of aberrantly glycosylated MUC1, which has shown encouraging activity in early clinical evaluation and further supports the therapeutic potential of this antigen class (31–36). Alternatively, CAR-T cell approaches(37–39), including 5E5-based constructs targeting the Tn-bearing GSTA epitope of MUC1, have likewise demonstrated potent glycoform-selective cytotoxicity and tumor control in preclinical models and have entered early-phase clinical evaluation. However, as with other CAR-T strategies for solid tumors, their activity may be constrained by limited persistence, tumor infiltration, and T cell dysfunction within the tumor microenvironment (39–42). Despite these advances, intratumoral glycan heterogeneity remains insufficiently addressed. Because Tn and STn frequently coexist at variable abundance within the same tumor, therapies restricted to a single glycoform remain vulnerable to antigen-driven escape. A strategy that maintains recognition of both glycoforms while efficiently recruiting antitumor immunity could therefore provide an important advantage.

Bispecific T-cell engagers (TCEs) offer a particularly attractive framework for this purpose because they redirect endogenous polyclonal T cells against tumor cells in an MHC-independent manner without requiring ex vivo cell manufacturing (43,44). TCEs are clinically validated in hematologic malignancies and are increasingly being developed for solid tumors. However, to our knowledge, no TCE has yet exploited a tumor-associated glycopeptide antigen such as MUC1-Tn/STn as its tumor-targeting arm. This gap likely reflects the structural complexity and context dependence of glycopeptide antigens, together with the lack of antibodies engineered to achieve high-affinity recognition across heterogeneous Tn/STn glycoforms(4,5,40,45).

Here, we report the structure-guided directed evolution of 5E5, an anti-MUC1-Tn antibody, to generate affinity-matured variants with high-affinity dual recognition of MUC1-Tn and MUC1-STn. Using yeast surface display and the SAMPLER (Synthetic Antibody Mutagenesis strategy by multiple Point Loop Enrichments and Repertoires) platform, we identified variants with significantly enhanced recognition of both glycoforms across defined glycopeptides, engineered tumor cells, and primary human tumor tissues while retaining minimal reactivity toward normal tissues. Co-crystal structures define the molecular basis of affinity enhancement and identify HCDR3 residue 98 as a key determinant that fine-tunes relative Tn versus STn preference. When reformatted as bispecific TCEs, the affinity-matured variants display enhanced cytolytic potency, overcome mixed Tn/STn antigen heterogeneity, achieve durable tumor suppression in vivo where the parental BiTE is ineffective, and show superior activity in patient-derived tumor-like clusters. Together, these findings establish a structural and functional framework for engineering glycan-heterogeneity-resilient immunotherapies for solid tumors.

## Results

### Structure-guided affinity maturation of 5E5 enhances dual recognition of MUC1-Tn and MUC1-STn

Monoclonal antibody 5E5 specifically recognize the GSTA region of the MUC1 VNTR upon O-GalNAcylation of the threonine residue(46,47). While parental 5E5 in scFv-Fc format possesses sub-micromolar baseline affinity for the MUC1-Tn glycoform, its cross-reactivity toward the sialylated derivative, MUC1-STn, is approximately 5-fold weaker (**Supplementary Fig S1A**). To define the molecular basis of MUC1-Tn recognition and guide affinity maturation, we analyzed the co-crystal structure of the wild-type (WT) 5E5 single chain variable fragment (scFv) bound to the MUC1-Tn glycopeptide GST(Tn)APPA (Ag_Tn), determined at 2.35 Å resolution (**Fig. 1A** and **Supplementary Figs S1 B-D, Supplementary table S1**).

**Figure 1.**
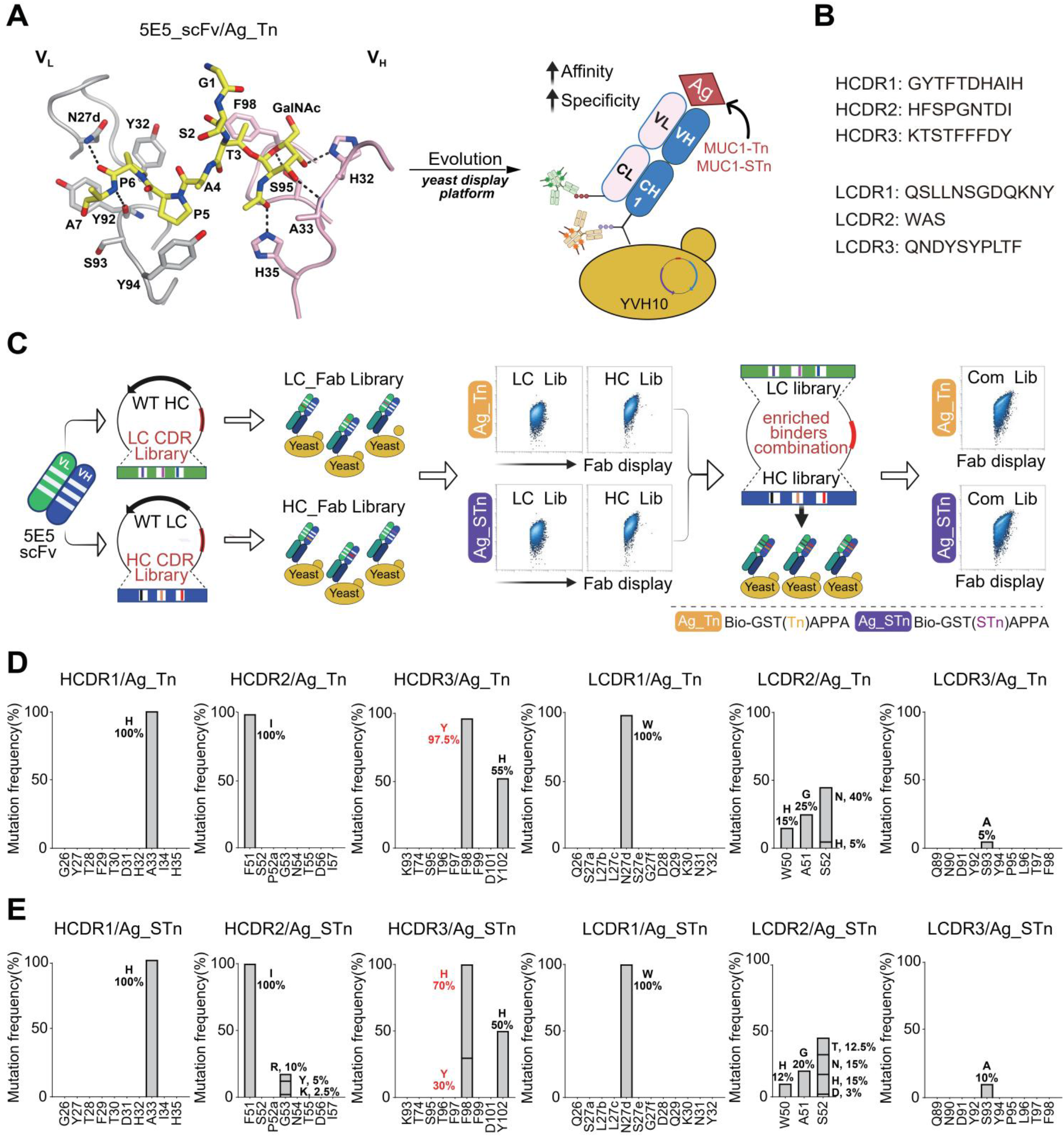
Structure-guided directed evolution strategy for affinity maturation of 5E5 against MUC1-Tn and MUC1-STn glycoepitopes. **A)** Schematic representation illustrates the glycoepitope recognition interface from the co-crystal structure of the 5E5 WT scFv in complex with the MUC1-Tn glycopeptide (Ag_Tn) as the template for evolving affinity and specificity towards both MUC1-Tn and MUC1-STn antigens via yeast display directed evolution on the 5E5 scaffold. Antibody light chain (grey) and heavy chain (light pink) are shown with Ag_Tn peptide backbone and GalNAc moiety displayed as yellow sticks. Black dashed lines indicate direct hydrogen bonds between antibody key residues (H32, A33, H35, S95, N27d, Y92) and Ag_Tn that define the glycoepitope recognition interface. **B)** CDR loop sequences of parental WT 5E5 used as the mutagenesis scaffold, showing all six complementarity-determining regions (HCDR1-3, LCDR1-3). **C)** Schematic overview of the SAMPLER (Synthetic Antibody Mutagenesis strategy by multiple Point Loop Enrichments and Repertoires) library construction and selection workflow. Two independent diversity libraries were constructed from the 5E5 scFv scaffold: an LC CDR library paired with WT HC, and an HC CDR library paired with WT LC. Each library was displayed on the yeast cell surface in Fab format and subjected to four rounds of FACS against biotinylated MUC1-Tn (Ag_Tn; Bio-GST(Tn)APPA) or MUC1-STn (Ag_STn; Bio-GST(STn)APPA). Enriched binders from HC and LC libraries were combined to generate a combinatorial library (Com Lib), which underwent an identical four-round FACS selection procedure to identify affinity-matured variants with optimized binding affinity, glycoform specificity, and conformational stability. Representative FACS dot plots show progressive enrichment of high-binding populations across selection rounds for both LC and HC libraries against each antigen (**HC**: heavy chain; **LC**: light chain; Ag: antigen; **Bio**: Biotin). **D)** Deep sequencing analysis of enriched binders from the combinatorial HC/LC library after four rounds of FACS selection against biotinylated Ag_Tn (Bio-GST(Tn)APPA) revealed recurrent mutations across all six CDR loops (HCDR1-3, LCDR1-3). Primary enrichment was observed at HCDR1 position 33 (His, 100%), HCDR2 position 51 (Ile, 100%), HCDR3 positions 98 (Tyr, 97.5%) and 102 (His, ∼55%), and LCDR1 position 27d (Trp, 100%). Secondary enrichment occurred at LCDR2 positions 50 (His, 15%), 51 (Gly, 25%), and 52 (Asn, 40%; His, 5%), with minor enrichment at LCDR3 position 93 (Ala, 5%) **E)** Deep sequencing analysis of enriched binders from the combinatorial HC/LC library after four rounds of FACS selection against biotinylated Ag_STn (Bio-GST(STn)APPA) revealed enrichment patterns largely consistent with Ag_Tn selections (bar charts presented as in **D**). Notably, HCDR3 position 98 showed differential enrichment: histidine (70%) was preferentially enriched over tyrosine (30%) in the Ag_STn condition. HCDR3 position 102 (His, ∼50%) was co-enriched with position 98 under both selection conditions. Secondary enrichment occurred at LCDR2 positions 50 (His, 12.5%), 51 (Gly, 20%), and 52 (Thr, 12.5%; Asn, 15%; His, 15%; Asp, 2.5%), with minor enrichment at LCDR3 position 93 (Ala, 10%).

Compared to the MUC1-Tn peptide APGST(Tn)AP in the previously reported co-crystal structure with WT 5E5 scFv at 3.0 Å resolution(47), the peptide Ag_Tn lacks the two N-terminal residues Ala-Pro (AP), which make no contact with antibody 5E5 and incorporates two C-terminal residues Pro-Ala (PA) that are critical for antibody binding (**Fig. 1A**). The co-crystal structure of Ag_Tn with 5E5 scFv exhibited ordered electron density for all residues and the GalNAc moiety (**Supplementary Fig S1D**). Approximately 310 Å^2^ and 200 Å^2^ of molecular surface are buried on the residues and the GalNAc moiety of Ag_Tn, respectively, while the 5E5 light (L) and heavy (H) chains contribute approximately 46% and 54% of the total peptide buried surface area, respectively. The GalNAc moiety, recognized by the 5E5 H-chain, is the critical component of Ag_Tn, contributing the majority of hydrogen bond interactions. The Ag_Tn C-terminal residues Pro6 and Ala7 contribute two hydrogen bonds to the 5E5 L-chain through their mainchain atoms, and Ag_Tn Pro5 and Pro6 engage with L-chain aromatic residue Tyr94 and Tyr32 through CH-π stacking interactions, respectively. The N-terminal residues Gly1 and Ser2 of Ag_Tn project out of the antigen-binding site groove and make no contacts with the antibody. HCDR1 and HCDR3 mediate direct contacts with the GalNAc moiety and the glycosylated Thr3, while LCDR1 and LCDR3 primarily interact with the four C-terminal residues Ala4, Pro5, Pro6 and Ala7 (**Fig. 1A**).

Guided by this structural information, we used SAMPLER(48), a yeast-display-based platform for synthetic antibody mutagenesis, to affinity-mature 5E5 (**Fig. 1A**). Two independent diversity libraries were constructed on the 5E5 scFv scaffold. In the light-chain CDR library, LCDR1–LCDR3 were diversified while the parental heavy chain was retained. Conversely, in the heavy-chain CDR library, HCDR1–HCDR3 were diversified while the parental light chain was retained. Each variant contained up to three point mutations. Library design focused primarily on CDR1 and CDR3, where each residue was diversified to sample all 20 amino acids. Although HCDR2 and LCDR2 do not directly contact Ag_Tn in the parental 5E5 structure, HCDR2 was fully diversified to explore mutations that could tune neighboring CDR conformations, reshape the GalNAc-binding pocket, or improve accommodation of the extended STn glycan. By contrast, only selected LCDR2 residues were sampled, allowing potential secondary contributions from the light chain to be captured while maintaining sufficient library coverage for yeast-display screening (**Fig. 1B**).

Both libraries were displayed on *Saccharomyces cerevisiae* in Fab format and subjected to four sequential rounds of fluorescence-activated cell sorting (FACS) using biotinylated MUC1-Tn glycopeptide (Ag_Tn; Bio-GST(Tn)APPA) or biotinylated MUC1-STn glycopeptide (Ag_STn; Bio-GST(STn)APPA). Rounds 1, 2, and 4 used titrated antigen concentrations to enrich high-binding populations, corresponding to the top 0.1%–15% of antigen-positive cells, whereas round 3 incorporated negative selection against biotinylated HEK293 membrane proteins to deplete nonspecific binders. Enriched heavy-and light-chain libraries were then recombined to generate a combinatorial HC/LC library, which was subjected to an identical four-round selection against both Ag_Tn and Ag_STn (**Fig. 1C, Supplementary Figs S2-S3**).

Deep sequencing of the enriched combinatorial libraries revealed a convergent mutational signature across both single-chain and combinatorial selections (**Figs. 1D-1E**). In the heavy chain, several positions showed strong enrichment, with both shared and glycoform-dependent patterns across the two selection conditions. The mutation Ala33His in HCDR1 and Phe51Ile in HCDR2 reached 100% frequency. By contrast, HCDR3 position 98 displayed a striking antigen-dependent divergence: Tyr98 was enriched to greater than 97.5% frequency under Ag_Tn selection, whereas His98 was preferentially enriched to approximately 70% frequency under Ag_STn selection, with Tyr98 remaining substantially represented at approximately 30%. Thus, residue 98 emerged as a key determinant that tunes glycoform preference between MUC1-Tn and MUC1-STn. A second HCDR3 substitution, His102, co-enriched with both Tyr98-and His98-containing variants, reaching approximately 55% and 50% frequency under Ag_Tn and Ag_STn selection, respectively. Because this substitution exceeded the nominal three-mutation design limit of the SAMPLER library, its recurrent enrichment suggests an independent contribution to antigen engagement.

In the light chain, LCDR1 carried the dominant enrichment signal, with N27dW reaching 100% frequency under both selection conditions. LCDR2 showed only modest enrichment, consistent with its limited structural engagement in the parental complex, whereas LCDR3 showed no appreciable enrichment. Together, these data show that directed evolution of 5E5 converges on a discrete set of mutations across HCDR1–HCDR3 and LCDR1–LCDR2, with HCDR3 residue 98 serving as the principal determinant of Tn *versus* STn glycoepitope preference.

### Affinity-matured 5E5 variants exhibit enhanced and selective recognition of MUC1-Tn/STn glycopeptides

Guided by the sequencing results and the dominant contribution of the heavy chain to glycan recognition, we selected eight representative affinity-matured variants (mut1–mut8) for biochemical and functional characterization. These variants were stratified according to their mutational patterns: mut1–mut4 contain six mutations distributed across the heavy-and light-chain CDRs, mut5–mut6 contain four HCDR mutations, whereas mut7–mut8 contain three (**Supplementary Fig S4A**). All variants share H33, I51, and H102 substitutions in the HCDR loops and carry either Tyr or His at HCDR3 position 98, defining two glycoform-preference groups. Mut1–mut4 additionally share W27d in LCDR1 and carry either H50 or N52 in LCDR2. All variants were produced as Fc fusions, with mut1–mut4 also generated as Fabs for kinetic characterization (**Fig. 2A** and (**Supplementary Fig S4A**).

**Figure 2.**
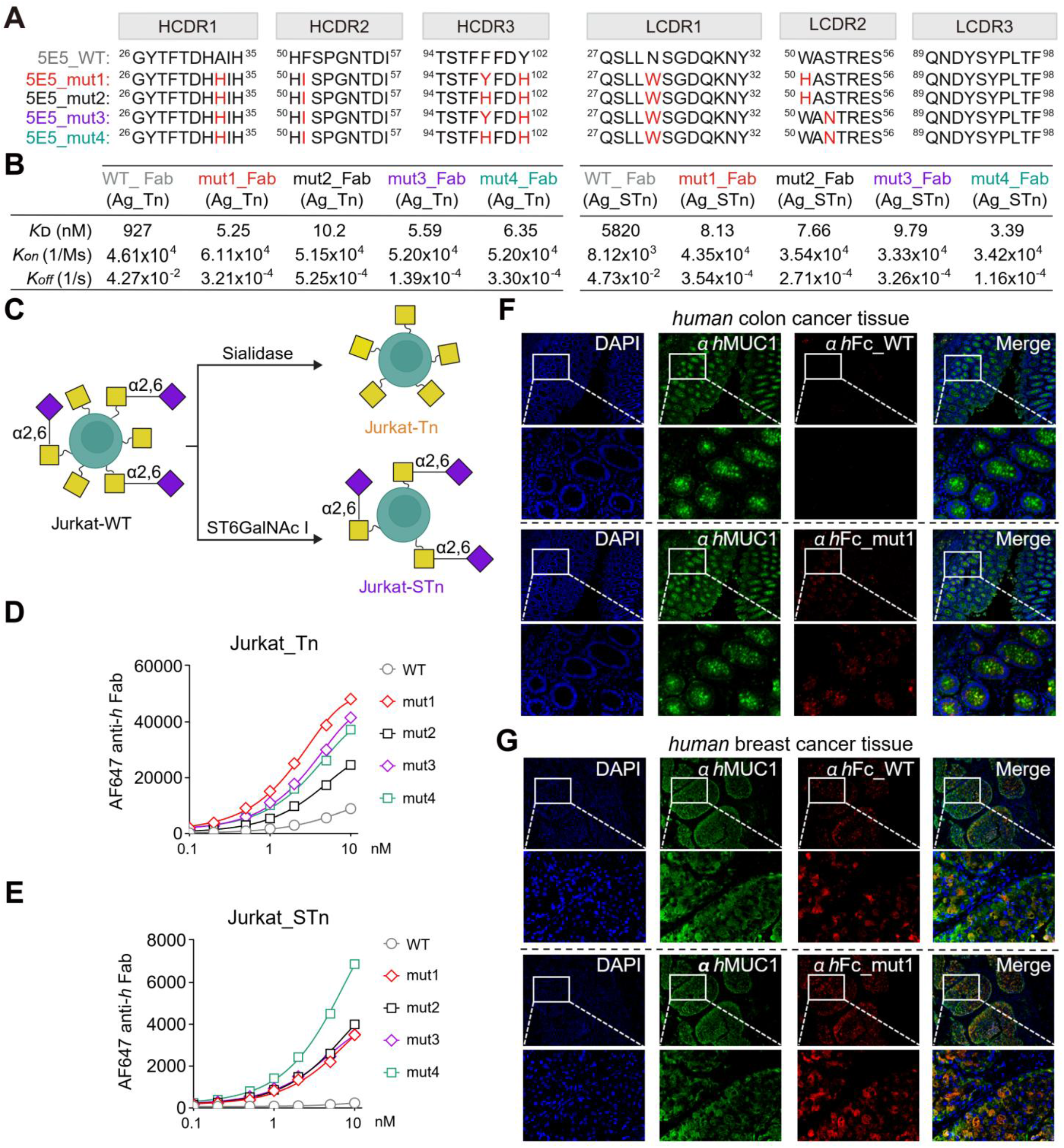
Enhanced recognition of MUC1–Tn and MUC1–STn peptides, glycoengineered cell lines and human tissues by affinity-matured 5E5 variants. **A)** CDR loop sequence alignment of wild-type 5E5 (5E5_WT) and four representative affinity-matured variants (5E5_mut1– 4). Mutated residues relative to wild-type are highlighted in red for each variant. Superscript numbers indicate Kabat residue positions. Key mutations include A^33^H (HCDR1), F^51^I (HCDR2), F^98^Y/H and Y^102^H (HCDR3), N^27d^W (LCDR1), and W^50^H or S^52^N (LCDR2). **B)** Binding kinetics of wild-type 5E5 Fab and affinity-matured mut1–4 Fab fragments toward Ag_Tn (Bio-GST(Tn)APPA, **left**) and Ag_STn (Bio-GST(STn)APPA, **right**) were determined by biolayer interferometry (BLI) using an Octet system. Equilibrium dissociation constants (*K*_D_), association rate constants (*k_on_*), and dissociation rate constants (*k_off_*) are presented. **C)** Schematic representation of glycosylation pattern engineering on Jurkat cells. Parental Jurkat_WT cells were treated with sialidase to generate Jurkat_Tn cells, or treated with sialyltransferase ST6GalNAc I to generate Jurkat_STn cells. **D, E)** Concentration-dependent binding of wild-type 5E5 Fab and affinity-matured mut1–mut4 Fab fragments to Jurkat_Tn cells (**D**) and Jurkat_STn cells (**E**), measured by flow cytometry and quantified as AF647 anti-human Fab mean fluorescence intensity (MFI) across a concentration range of 0.1–10 nM. **F)** Immunofluorescence staining of human colon cancer tissue sections with WT 5E5-Fc (top two rows) and mut1-Fc (bottom two rows). Sections were co-stained with anti-human MUC1 antibody (αhMUC1, green) as a reference for endogenous MUC1 expression, and DAPI (blue) for nuclear staining. Anti-human Fc antibody detection (red) reveals binding of WT-Fc and mut1-Fc to MUC1-Tn/STn glycoepitopes within tumor tissue. Merge panels show overlay of all channels. Scale bar, 100 μm. **G)** Immunofluorescence staining of human breast cancer tissue sections with WT 5E5-Fc (top two rows) and mut1-Fc (bottom two rows), as in (**F**). Scale bar, 100 μm.

ELISA profiling of Fc-fusion variants against MUC1-Tn and MUC1-STn glycopeptides revealed substantially enhanced binding relative to parental 5E5, with pronounced leftward shifts in the dose-response curves. A clear binding hierarchy emerged across variants with increasing numbers of enriched mutations: mut1–mut4, incorporating six mutations across both heavy and light chains, achieved half-maximal binding at substantially lower concentrations than parental 5E5 (Ag_Tn: WT EC_50_ = 37.3 nM *versus* mut1–mut4 EC_50_ = 0.10–0.18 nM; Ag_STn: WT EC_50_ = 185 nM *versus* mut1–mut4 EC_50_ = 0.26–0.88 nM), whereas mut5–mut6 and mut7–mut8 showed progressively weaker binding as the number of incorporated HCDR mutations decreased from four to three. Consistent with the selection profiles, position 98 also influenced relative glycoform preference. Tyr98-containing variants consistently showed lower EC_50_ values for Ag_Tn across multiple variant pairs (mut1 *vs* mut2, 0.10 *vs* 0.18 nM; mut5 *vs* mut6, 0.19 *vs* 0.43 nM; mut7 *vs* mut8, 0.36 *vs* 0.88 nM), whereas His98-containing variants showed lower EC_50_ values for Ag_STn (mut1 *vs* mut2, 0.56 *vs* 0.26 nM; mut3 *vs* mut4, 0.88 *vs* 0.31 nM) (**Supplementary Figs S4B–S4E**)

Biolayer interferometry (BLI) analysis of mut1–mut4 Fab fragments further defined the kinetic basis of affinity enhancement (**Figs. 2A-2B** and **Supplementary Figs S4F-S4O**). For MUC1-Tn, mut1–mut4 achieved approximately 90-to 177-fold improvements in affinity relative to parental 5E5 Fab, lowering the *K*_D_ from the micromolar to nanomolar range (927 nM to 5.25–10.2 nM). These improvements were driven primarily by substantially reduced dissociation rates, with largely preserved association rates. Affinity gains for MUC1-STn were even more pronounced, reaching approximately 600-to 1700-fold improvements in affinty (5820 nM to 3.39–9.79 nM), reflecting both faster association and slower dissociation. BLI measurements further substantiated the contribution of position 98 to relative glycoform preference: mut2 (His98, *K*_D_ = 10.2) showed approximately 2-fold weaker Tn affinity than mut1 (*K*_D_ 5.25 nM), whilst mut4 (His98) achieved the highest STn affinity among all variants (*K*_D_ = 3.39 nM), approximately 2.9-fold better than mut3 (*K*_D_ = 9.79 nM). Importantly, all four variants exhibited high-affinity binding to both glycoforms, demonstrating that directed evolution strengthened overall antigen engagement across both MUC1-Tn and MUC1-STn.

We next assessed epitope specificity using a panel of six antigens comprising paired unglycosylated and Tn-glycosylated MUC1 peptides spanning the GVTS, PDTR, and GSTA epitope regions (**Supplementary Figs S5A-S5B**). Mut1–mut4 Fc-fusion variants selectively recognized GST(Tn)APPA, with no detectable binding to the unglycosylated GSTAPPA control, confirming strict dependence on Tn glycosylation for recognition. No detectable binding to GVTS-or PDTR-region peptides in either their glycosylated or unglycosylated forms **(Supplementary Fig S5C**). Thus, affinity maturation preserves the glycan-dependent epitope selectivity of parental 5E5 while substantially enhancing binding to its target MUC1 glycoepitopes.

### Affinity-matured 5E5 variants bind cancer cell lines and human tumor tissues with high specificity

We then evaluated the affinity-matured variants in cell-surface binding assays using Jurkat cells engineered to display either Tn or STn antigens. Jurkat_Tn cells were generated by sialidase treatment, whereas Jurkat_STn cells were produced through ST6GalNAc I-catalyzed α2,6-sialylation (**Fig. 2C**). All four affinity-matured variants, mut1–mut4, bound both Jurkat_Tn and Jurkat_STn cells with substantially higher affinity than parental 5E5, and the Tyr98/His98-dependent preference observed in biochemical assays was retained at the cellular level (**Figs. 2D**-2**E**). By contrast, all variants showed negligible binding to MUC1-negative HEK293 cells (**Supplementary Fig S5D**), indicating minimal nonspecific cell-surface engagement.

Because therapeutic targeting of MUC1-Tn/STn requires not only enhanced glycoepitope binding but also stringent tumor selectivity, we next evaluated the binding of Fc-fused variants using a panel of human tumor and normal tissues by Immunofluorescence staining. Mut1 Fc-fusion protein showed stronger staining than parental 5E5 in MUC1-Tn/STn-positive regions of colon and breast cancer tissues, with prominent colocalization with anti-MUC1 reference staining confirming on-target tumor engagement (**Figs. 2F**-**2G**). Notably, mut1 showed minimal cross-reactivity across a panel of six normal human tissues, including lung, placenta, colon, skin, adrenal gland, and tonsil (**Supplementary Fig S5E**). Together, these results demonstrate that affinity maturation enhances tumor-associated MUC1-Tn/STn recognition while preserving the antigen and tissue selectivity required for therapeutic translation.

### Structural basis of affinity maturation and glycoform preference

To define the atomic basis of affinity maturation and the glycoform preference at HCDR3 position 98, we determined co-crystal structures of mut2 Fab and mut3 Fab bound to Ag_Tn at 1.59 Å and 1.66 Å resolution, respectively, as well as mut3 Fab bound to Ag_STn at 1.42 Å resolution (**Figs. 3A–3C**, **Supplementary Fig S6**, **Table S1**). Relative to WT 5E5, mut2 and mut3 each contain six mutations distributed across all CDR loops except LCDR3 (**Supplementary Fig S7**).

**Figure 3.**
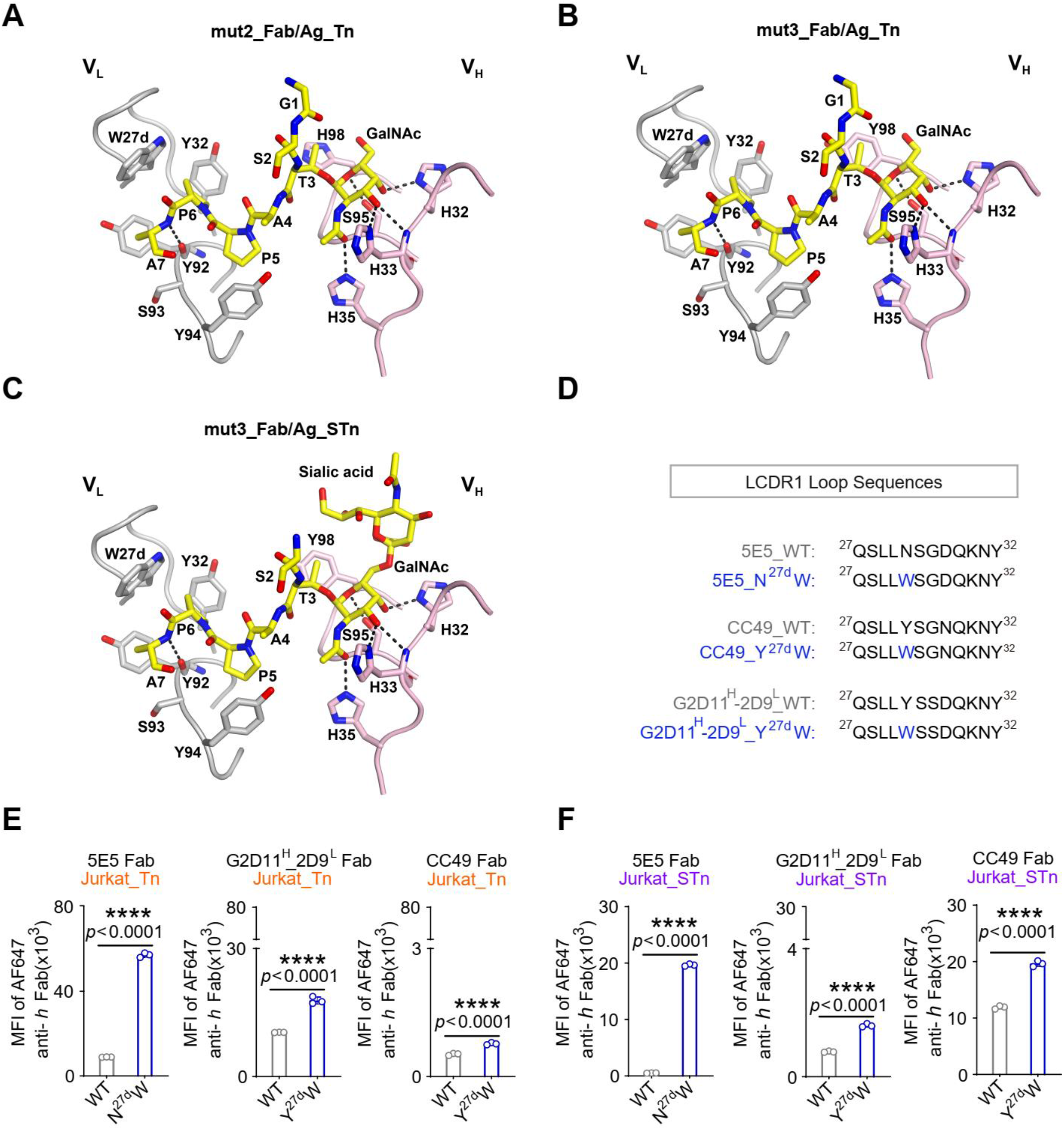
Structural characterization of MUC1-Tn and MUC1-STn recognition by parental and affinity-matured 5E5 antibodies and transferability of affinity-maturation mutations to distinct anti-MUC1-Tn/STn antibodies. **A)** Co-crystal structure of mut2 Fab in complex with MUC1-Tn glycopeptide (Ag_Tn). Antibody light chain (L-chain, grey) and heavy chain (H-chain, light pink) are shown with the glycopeptide carbon atoms displayed as yellow sticks with nitrogens in blue and oxygens in red. **B)** Co-crystal structure of mut3 Fab in complex with Ag_Tn, displayed as in (**A**). **C)** Co-crystal structure of mut3 Fab in complex with MUC1-STn glycopeptide (Ag_STn). Antibody light chain and heavy chain are shown as in (**A**-**B**), with the glycopeptide and sialic acid moiety displayed as yellow sticks. **D)** LCDR1 loop sequences of wild-type and mutant variants of three anti-MUC1-Tn/STn antibodies: 5E5, chimeric G2D11^H^-2D9^L^ and CC49. The transferable N/Y²⁷ᵈW mutation at LCDR1 position 27d is highlighted in blue; WT residues are shown in black. Kabat numbering is used throughout. **E, F**) Flow cytometric binding analysis of wild-type and mutant Fab fragments from 5E5, G2D11-2D9 and CC49 antibody families against Jurkat_Tn cells (**E**) and Jurkat_STn cells (**F**). Data represent mean ± SEM of n = 3 independent experiments. Two-tailed unpaired Student’s *t*-test. ns, not significant; \**p* < 0.05; \*\**p* < 0.01; \*\*\**p* < 0.001; \*\*\*\**p <* 0.0001.

The Ag_Tn-bound mut2 and mut3 Fab structures were nearly identical, with an RMSD of 0.1 Å over the Cα atoms of the Fv domains, and both aligned closely onto the WT 5E5 scFv structure, with an RMSD of 0.6 Å. Ag_Tn occupied a binding pose similar to that observed in the WT 5E5 complex and retained the core recognition features, including hydrogen bonds between GalNAc and the heavy chain and CH-π stacking interactions between the C-terminal peptide residues Pro5 and Pro6 and the light chain residues Tyr94 and Tyr32, respectively (**Figs. 3A–3C**). Thus, affinity maturation does not remodel the overall binding mode of 5E5 but instead strengthens antigen engagement through local, mutation-driven interactions.

Mut2 and mut3 share four affinity-matured substitutions (**Supplementary Fig S7**). In LCDR1, Trp27d replaces the parental Asn27d and positions its indole side chain as an interfacial anchor against the hydrophobic C-terminal residues Pro6 and Ala7 of Ag_Tn or Ag_STn. Although this substitution eliminates a hydrogen bond between Asn27d and the backbone carbonyl of Pro6, it introduces a larger hydrophobic contact surface that likely stabilizes peptide engagement (**Figs. 1A** and **3A–3C**). In HCDR1, the mutation Ala33His creates an additional hydrogen bond between the His33 side chain imidazole and the O3 hydroxyl of GalNAc in Ag_Tn or Ag_STn (**Figs. 3A– 3C**). This substitution is structurally coupled to the HCDR2 Phe51Ile mutation. The bulkier His33 side chain shifts HCDR2 residues 52–56 away from the peptide, displacing the Cα atom of Asn54 by 2.6–2.7 Å in mut2 and mut3 Fabs; replacement of Phe51 with Ile reduces local steric bulk and avoids a potential clash with Thr55 (**Supplementary Fig S8**). Thus, Ala33His and Phe51Ile act cooperatively to accommodate HCDR2 remodeling while preserving the geometry of the antigen-binding site.

The shared HCDR3 Tyr102His mutation is positioned distal to the glycopeptide but may fine-tune an intra-domain aromatic network that supports GalNAc recognition. In WT 5E5, Tyr102, Phe99, and His32 form an aromatic interaction network in which Tyr102 engages Phe99 through a T-shaped aromatic interaction, while Phe99 stacks with HCDR1 His32. His32 also contacts GalNAc directly, including through a hydrogen bond with the O4 hydroxyl group (**Supplementary Fig S9**). This network is preserved in mut2 and mut3, with the His102 imidazole maintaining aromatic interactions with Phe99. Although Tyr102His does not directly contact antigen, its recurrent enrichment suggests that it may modulate the geometry or stability of this GalNAc-engaging network.

Mut2 and mut3 differ at three affinity-matured positions, most notably HCDR3 residue 98. In WT 5E5, Phe98 forms part of the binding-groove wall and contacts both GalNAc and the glycosylated Thr3. In mut2 and mut3, Phe98 is replaced by His98 and Tyr98, respectively. Both side chains occupy a position similar to WT Phe98 and retain contacts with GalNAc and Thr3 in Ag_Tn or Ag_STn. Notably, His98 and Tyr98 each form a water-mediated hydrogen bond to the backbone carbonyl of Thr3 through two bridging water molecules (**Supplementary Fig S10**), providing a structural basis for the enhanced engagement of the glycopeptide core.

Mut2 and mut3 also differ at LCDR2 positions 50 and 52, which are distal to the peptide but connect structurally to HCDR3 residue 98. In mut2, LCDR2 His50 forms a water-mediated hydrogen-bonding network with HCDR3 His98 through two bridging water molecules. In mut3, LCDR2 Asn52 hydrogen bonds to Thr53 that interacts with LCDR2 Trp50, which in turn contacts HCDR3 Tyr98 (**Supplementary Fig S11**). These distal LCDR2 substitutions therefore appear to support the local conformation of the HCDR3 selectivity residue rather than directly contacting antigen.

Comparison of the mut3 Fab structure bound to Ag_Tn and Ag_STn further clarified the basis of glycoform preference (**Figs. 3B-3C**). Both glycopeptides adopt similar binding poses, with comparable conformations of the peptide backbone and GalNAc moiety, although Gly1 of Ag_STn was not modeled because of weak electron density. The sialic acid moiety of Ag_STn projects outward from the binding pocket and makes only limited contacts with the antibody, primarily van der Waals interactions with HCDR1 His32 and HCDR3 Tyr98. These structural observations provide a basis for the enhanced affinity of both His98-and Tyr98-bearing variants through additional water-mediated contacts with the Thr3 backbone carbonyl, whilst the differential glycoform preference observed kinetically likely reflects subtle differences in sialic acid accommodation not fully resolved in the current structures.

We next asked whether the structural principles revealed by 5E5 affinity maturation could be used to guide engineering of related anti-Tn/STn antibodies. Several antibodies against tumor-associated Tn/STn glycoepitopes have been reported, including G2D11(49,50), 2D9(50,51), and CC49(52). G2D11 recognizes adjacent Tn antigens in di-Tn motifs with comparable specificity across Ser-Thr, Thr-Thr, and Thr-Ser glycosylation patterns, and its binding is minimally influenced by the surrounding peptide sequence, reflecting a predominantly glycan-driven mode of recognition(49). By contrast, 2D9 binds the same GSTAPP region of the MUC1 VNTR as 5E5 but has distinct epitope requirements: whereas 5E5 recognizes the minimal GST(Tn)AP epitope, 2D9 binds to the di-glycosylated GS(Tn)T(Tn)AP motif (46,51). CC49 is a well-characterized antibody targeting the STn-bearing TAG-72 antigen and has been extensively evaluated for tumor targeting. Structural comparison with 5E5 reveals a shared architecture in which the heavy chain primarily engages GalNAc, whereas the light chain contacts the underlying peptide scaffold(50,52). Sequence alignments further identified conserved features in HCDR1, HCDR2, and LCDR1 overlapping with regions that acquired convergent mutations during 5E5 affinity maturation. These similarities raised the possibility that selective affinity-enhancing mutations identified in 5E5 could be transferred to related anti-glycopeptide antibody frameworks(50).

We reasoned, however, that the heavy-chain mutations would be difficult to transfer directly because they act cooperatively to shape the GalNAc-binding site, and HCDR3 differs substantially in both sequence and length across antibody families. We therefore focused on the convergently enriched LCDR1 W27d substitution, which strengthens peptide engagement. The equivalent Y27dW substitution was introduced into CC49 and chimeric G2D11^H^-2D9^L^, respectively, and an N27dW single mutant was generated in 5E5 for comparison (**Fig 3D**).

Binding assays using Jurkat_Tn and Jurkat_STn cells showed that W27d substitution significantly enhanced recognition of both glycoforms across all three antibody scaffolds relative to their corresponding wild-type antibodies (**Figs. 3E-3F**). Although the magnitude of improvement was scaffold-dependent and greatest in 5E5, consistent gains were observed in CC49 and G2D11^H^-2D9^L^, indicating a transferable mechanism across related anti-glycopeptide scaffolds. Importantly, W27d introduction did not increase non-specific binding to MUC1-negative HEK293T cells in any scaffold (**Supplementary Fig S12**), confirming that the enhanced cell-surface engagement is strictly antigen-dependent. Together, these results establish W27d as a generalizable affinity-enhancing substitution for engineering antibodies against tumor-associated MUC1 glycoforms.

### Affinity-matured 5E5 BiTE variants exhibit enhanced cytolytic activity

To determine whether enhanced MUC1-Tn/STn recognition translates into improved T-cell-mediated cytotoxicity, we reformatted mut1 and mut2 as bispecific T-cell engagers (BiTEs). Each BiTE contained an affinity-matured 5E5 Fab as the tumor-targeting arm and an anti-CD3 scFv as the T-cell-engaging arm, joined by a flexible (G₄S)n linker (**Fig. 4A**). We first generated three mut1-BiTE formats with increasing linker length (1L, 2L, and 3L) to optimize molecular geometry. All three formats bound MUC1-Tn/STn-expressing MCF7 cells and human PBMCs comparably, with markedly enhanced tumor-cell binding relative to WT-BiTE but equivalent PBMC binding, indicating that affinity maturation selectively improved tumor antigen recognition without altering CD3 engagement (**Supplementary Figs S13A-S13B**). Consistent with this binding profile, all three mut1-BiTE formats induced stronger tumor-cell-dependent CD25^+^ upregulation in CD3^+^ T cells and mediated superior tumor-cell killing compared with WT-BiTE, with no significant differences among linker lengths (**Supplementary Figs S13C-S13E**). The single-linker G₄S format was therefore selected for subsequent studies.

**Figure 4.**
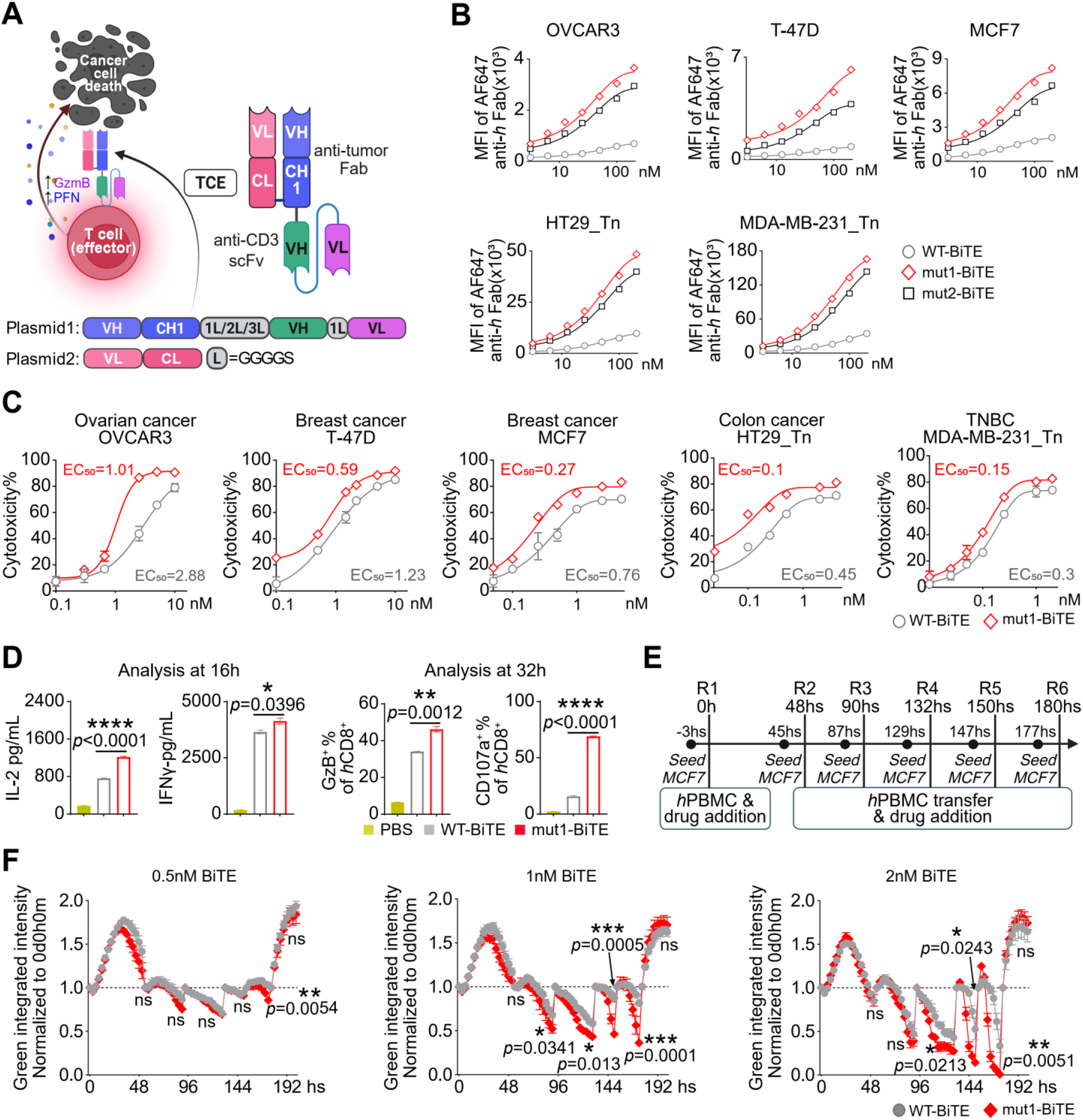
Affinity-matured 5E5 BiTE variants trigger enhanced tumor cell targeting, cytotoxicity and long-term killing potency against MUC1-Tn/STn-expressing tumors. **A)** Schematic representation of the bispecific T cell engager (BiTE) construct format and functional mechanism. The BiTE comprises an anti-tumor Fab arm targeting MUC1-Tn/STn glycoepitopes connected via a flexible (Gly₄Ser)n linker to an anti-CD3 scFv that engages CD3ε on cytotoxic T lymphocytes. BiTE-mediated crosslinking forms an immunological synapse driving T cell activation, granzyme B (GzmB) and perforin (PFN) secretion, and tumor cell death. Two plasmids were designed for encoding the heavy chain of anti-tumor Fab fused anti-human CD3 scFv (Plasmid 1: VH–CH1(anti-tumor)– 1L–VH (anti CD3)) and light chain of anti-tumor (Plasmid 2: VL–CL). L = (Gly₄Ser) linker unit; 1L/2L/3L indicates linker length variants tested. **B)** Flow cytometric binding analysis of WT-BiTE, mut1-BiTE, and mut2-BiTE across five MUC1-Tn/STn-positive cancer cell lines: OVCAR3 (ovarian cancer), T-47D (breast cancer), MCF7 (breast cancer), HT29_Tn (colon cancer, Tn-enriched), and MDA-MB-231_Tn (triple-negative breast cancer, Tn-enriched). Mean fluorescence intensity (MFI) of AF647-conjugated anti-human Fab is plotted against BiTE concentration (1–200 nM). **C)** IncuCyte-based cytotoxicity assays comparing WT-BiTE (grey) and mut1-BiTE (red) across five cancer cell lines: OVCAR3, T-47D, MCF7, HT29_Tn, and MDA-MB-231_Tn. Cytotoxicity (%) is plotted against BiTE concentration (0.1–10 nM). Human PBMCs were used as effector cells at a 5:1 effector-to-target (E:T) ratio.EC_50_ values for each condition are indicated. Data represent mean ± SEM of n = 3 independent experiments. **D)** Assessment of T cell cytokine production and effector molecule expression. IL-2 production (pg/mL), IFN-γ production (pg/mL), granzyme B (GzB) expression (% of CD8^+^ T cells), and CD107a (LAMP-1) surface expression (% of CD8^+^ T cells) measured at 16 h and 32 h following exposure to PBS, WT-BiTE, or mut1-BiTE. Measurements performed by ELISA (IL-2 and IFN-γ) and flow cytometry (GzB and CD107a). Data represent mean ± SEM of n = 3 independent experiments. Two-tailed unpaired Student’s t-test: *p < 0.05; **p < 0.01; ****p < 0.0001. **E)** Schematic of the serial killing assay design. MCF7 target cells were seeded at −3 h of round 1 (R1) and human PBMCs plus BiTE were added at 0 h to initial first round of killing. Fresh MCF7 target cells were re-seeded at rounds 2–6 (45, 87, 129, 147 h and 177h) with concurrent human PBMC transfer and BiTE re-addition. Normalized green integrated intensity was used to quantify surviving target cells at each time point. **F)** Serial killing assay results for WT-BiTE and mut1-BiTE at three concentrations: 0.5 nM (**left**), 1 nM (**middle**), and 2 nM (**right**). Normalized green integrated intensity is plotted over time (hours) for WT-BiTE (grey) and mut1-BiTE (red). Data are mean ± SEM of n = 3 independent experiments. Two-tailed unpaired Student’s *t*-test. ns, not significant; \**p* < 0.05; \*\**p* < 0.01; \*\*\**p* < 0.001; \*\*\*\**p <* 0.0001.

To evaluate tumor cell binding across a glycan-heterogeneous cancer cell line panel, dose-dependent binding of WT-BiTE, mut1-BiTE, and mut2-BiTE was assessed by flow cytometry in five MUC1-Tn/STn-expressing cancer cell lines including ovarian carcinoma OVCAR3, breast carcinoma T-47D and MCF7, colon cancer *COSMC-knockout* HT29-Tn and triple-negative breast cancer *COSMC-knockout* MDA-MB-231-Tn(7) (**Fig. 4B**). Both mut1-and mut2-BiTEs demonstrated markedly enhanced dose-dependent tumor cell binding relative to WT-BiTE across all five cell lines, with WT-BiTE showing essentially flat dose-response curves indicative of minimal engagement. The binding signal was highest in *COSMC-knockout* HT29_Tn and MDA-MB-231_Tn cells, which express the densest Tn epitopes, and progressively lower in wild-type cell lines with lower Tn density, consistent with an association between BiTE binding and surface MUC1-Tn/STn epitope abundance. Mut1-BiTE consistently showed superior binding over mut2-BiTE across all five cell lines. This pattern was consistent with the predominance of Tn over STn epitopes revealed by profiling with multiple anti-Tn/STn antibodies (**Supplementary Figs S13F-S13I**) and with the preferential Tn recognition associated with Tyr98 at HCDR3 position 98.

We then evaluated whether these binding gains translated into enhanced cytotoxic activity. In an IncuCyte-based killing assay across five target Tn-predominant cell lines, mut1-BiTE induced significantly stronger tumor-cell killing than WT-BiTE, with 2-to 5-fold reductions in EC_50_ and modest increases in maximal killing across two independent PBMC donors (**Fig. 4C** and **Supplementary Fig S14A**). These functional improvements were accompanied by increased CD25 and CD107a expression, granzyme B release, and IFN-γ and IL-2 secretion compared with WT-BiTE (**Fig. 4D**, **Supplementary Figs S13J, S15**). CD107a mobilization and granzyme B production were preferentially increased in CD8^+^ T cells relative to CD4^+^ T cells, indicating predominant engagement of cytotoxic T-cell function (**Supplementary Fig S15**). Together, these data suggest that affinity maturation primarily lowers the threshold for productive T-cell engagement rather than substantially increasing the ceiling of maximal cytotoxicity. This pattern is consistent with the reduced dissociation rates observed by BLI, which may prolong target engagement and facilitate productive T-cell activation. Serial killing assays further confirmed that mut1-BiTE sustained robust, dose-dependent cytotoxicity against MCF7 and T-47D cells over multiple rounds of sequential target-cell engagement, with consistently greater cytotoxic activity than WT-BiTE across the concentrations tested. (**Figs. 4E**-**4F**, **Supplementary Figs S14B**-**S14C**).

### Affinity-matured 5E5 BiTE variants suppress tumor growth in a heterogeneous Tn/STn model

The heterogeneous co-expression of Tn and STn glycoforms within individual tumors presents a challenge for glycan-targeted immunotherapy, as agents recognizing only one glycoform may provide incomplete coverage of heterogeneous tumor-cell populations. To determine whether affinity-matured dual-glycoform recognition could broaden functional coverage across heterogeneous tumor populations, we first examined the binding preferences of mut1-and mut2-BiTEs in STn-expressing cells. Binding evaluation on HT29-STn and MDA-MB-231-STn cells demonstrated that mut2-BiTE exhibited markedly stronger binding than both mut1-BiTE and WT-BiTE (**Figs. 5A-5B**). This pattern contrasted with the Tn-predominant cell lines, in which mut1-BiTE showed stronger binding (**Fig. 4B**), further supporting HCDR3 position 98 as a key determinant of relative glycoform preference.

**Figure 5.**
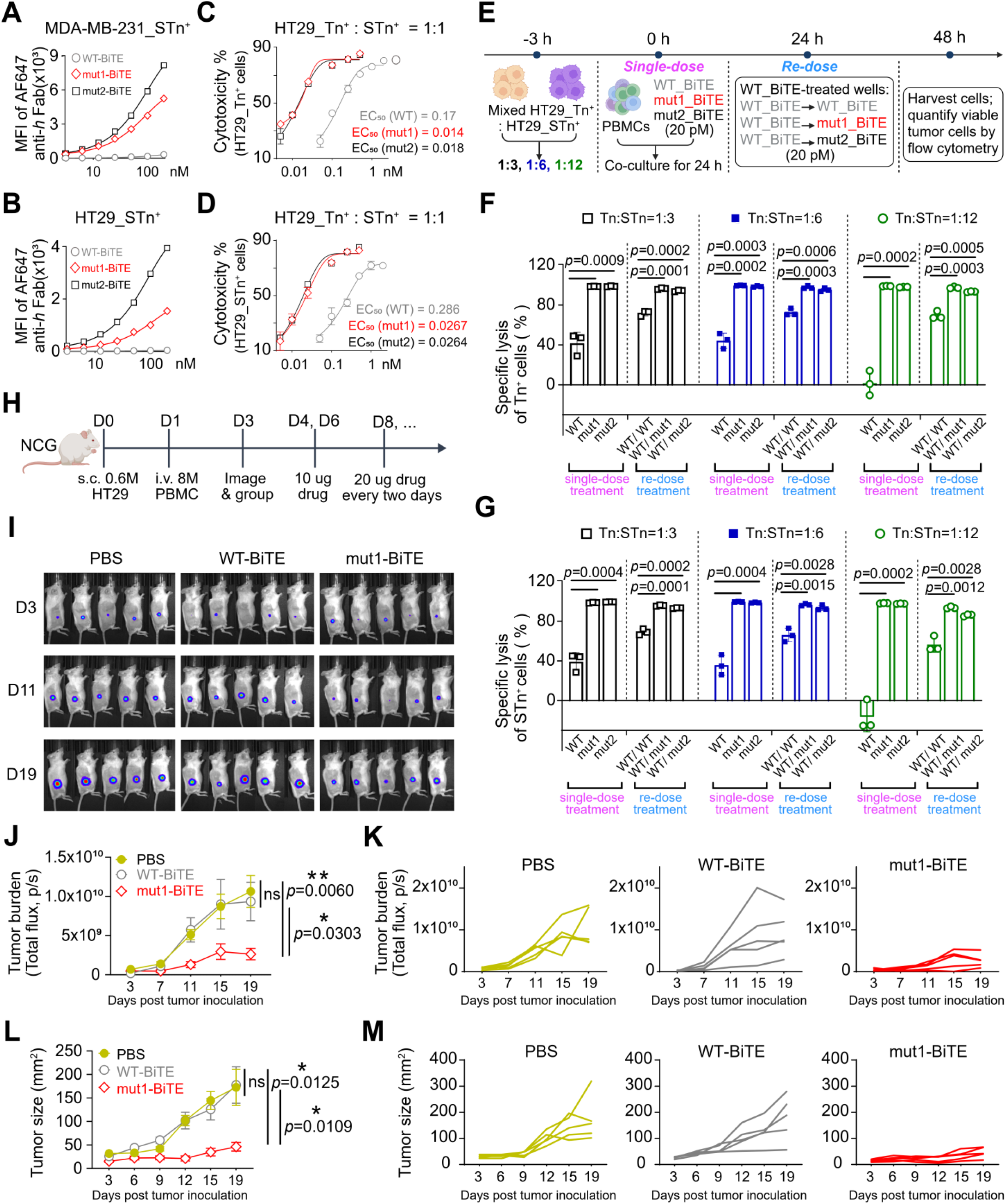
Affinity-matured BiTEs enhance antitumor activity in heterogeneous tumor models *in vitro* and *in vivo*. A, **B)** Flow cytometric binding analysis of WT-BiTE, mut1-BiTE, and mut2-BiTE to STn-high cancer cell lines MDA-MB-231_STn^+^ (**A**) and HT29_STn^+^ (**B**). Mean fluorescence intensity (MFI) of AF647-conjugated anti-human Fab is plotted against BiTE concentration (0.1–1000 nM). **C, D)** Dose-dependent cytotoxicity of WT-, mut1-, and mut2-BiTE against HT29_Tn⁺ (**C**) and HT29_STn⁺ (**D**) tumor cells in a mixed population at a Tn⁺: STn⁺ ratio of 1:1. EC_50_ values are indicated. **E)** Schematic of the single-dose and re-dose treatment experimental design. HT29_Tn⁺ and HT29_STn⁺ cells were mixed at the indicated ratios and co-cultured with PBMCs. At 0 h, cells were treated with WT-, mut1-, or mut2-BiTE (20 pM). After 24 h, WT-BiTE-treated wells were re-dosed with WT-, mut1-, or mut2-BiTE (20 pM). At 48 h, cells were harvested and viable tumor cells were quantified by flow cytometry. **F, G)** Specific lysis of HT29-Tn^+^ cells (**F**) and HT29-STn^+^ cells (**G**) following single-dose or re-dose BiTE treatment in mixed Tn^+^: STn^+^ co-culture. **H)** Schematic of the *in vivo* experimental design. Humanized NCG mice were inoculated subcutaneously with 0. 6×10^6^ cells comprising a 1:1 mixture of colon cancer cells HT29-Tn^+^ and HT29-STn^+^ on day 0, followed by engraftment with 6×10^6^ human PBMCs on day 1. Mice were imaged by bioluminescence on day 3 and randomized into three treatment groups PBS, WT-BiTE, and mut1-BiTE, receiving an initial dose of 10 μg on days 4 and 6, followed by 20 μg every two days thereafter. **I)** Representative bioluminescence images of tumor-bearing mice from the PBS, WT-BiTE, and mut1-BiTE groups at the indicated time points. **J)** Tumor burden quantified by bioluminescence imaging (total flux, photons/second) in PBS-, WT-BiTE-, and mut1-BiTE-treated mice over 19 days post tumor inoculation. **K)** Tumor burden trajectories of individual mice in each treatment group. **L)** Tumor size (mm²) measured by caliper in PBS-, WT-BiTE-, and mut1-BiTE-treated mice over 19 days post tumor inoculation. **(M)** Individual tumor growth curves in each treatment group. Data are presented as mean ± SEM. Statistical significance at Day 19 was determined by Student’s t-test; p values are indicated. Two-tailed unpaired Student’s *t*-test. ns, not significant; \**p* < 0.05; \*\**p* < 0.01; \*\*\**p* < 0.001; \*\*\*\**p <* 0.0001.

To evaluate cytolytic activity under conditions that more closely recapitulate intratumoral glycan heterogeneity, we first assessed BiTE killing efficacy in a 1:1 mixed HT29-Tn:HT29-STn co-culture model, in which Tn-and STn-expressing cells were present simultaneously (**Figs. 5C-5D**). Both mut1-and mut2-BiTEs demonstrated significantly enhanced killing of both Tn-and STn-expressing cells relative to WT-BiTE. For HT29-Tn cells, EC_50_ values decreased from 0.17 nM for WT-BiTE to 0.014 and 0.018 nM for mut1-and mut2-BiTE, respectively, corresponding to ∼12-and ∼9-fold improvements. For HT29-STn cells, EC_50_ values decreased from 0.29 nM for WT-BiTE to 0.0267 and 0.0264 nM for mut1-and mut2-BiTE, respectively, representing ∼11-fold improvements for both variants. These results demonstrate that affinity-matured BiTEs achieved potent cytolytic activity against both glycoforms simultaneously within a heterogeneous antigen environment.

To systematically evaluate the impact of intratumoral glycoform heterogeneity on BiTE efficacy, we established a model system in which mCherry-labeled HT29-Tn and GFP-labeled HT29-STn cells were combined at five defined Tn:STn ratios (6:1, 3:1, 1:1, 1:3, and 1:6) (**Supplementary Fig S16**). Under these heterogeneous conditions, mut1-and mut2-BiTEs induced greater T-cell degranulation than WT-BiTE across all Tn:STn ratios, as measured by CD107a mobilization in both CD8^+^ and CD4^+^ T cells (**Supplementary Figs S16A-S16B**). Consistently, both mut1-and mut2-BiTE achieved significantly greater cytotoxicity against both mCherry-labeled HT29-Tn and GFP-labeled HT29-STn cells than WT-BiTE across all five ratios (**Supplementary Figs S16C– S16E**). By contrast, the activity of WT-BiTE against STn^+^ cells progressively declined as the proportion of STn^+^ cells increased, highlighting the limited functional coverage of the parental BiTE for heterogeneous glycoform compositions.

We next asked whether affinity maturation could preserve cytotoxic activity under increasingly STn-dominant conditions, where WT-BiTE activity was most compromised. Mixed HT29-Tn and HT29-STn cultures were generated at Tn:STn ratios of 1:3, 1:6, and 1:12 and subjected to single-dose and re-dose killing assays. Cells were co-cultured with PBMCs and treated with BiTEs at 20 pM, a concentration near the EC_50_ values of mut1-and mut2-BiTEs against HT29-STn cells (**Fig. 5E**). Under these conditions, mut1-and mut2-BiTEs retained potent cytolytic activity against both Tn^+^ and STn^+^ tumor populations across all three ratios, whereas WT-BiTE activity progressively deteriorated as STn^+^ cells became more abundant. Notably, this loss of activity was observed not only against the STn^+^ population but also against the residual Tn^+^ cells, indicating that increasing STn dominance impaired overall WT-BiTE-mediated tumor clearance within the mixed population. At the most STn-dominant 1:12 ratio, WT-BiTE produced little to no cytotoxicity against either population, whereas mut1-and mut2-BiTEs maintained robust killing. To determine whether residual tumor cells escaping WT-BiTE treatment remained susceptible to the affinity-matured BiTEs, cultures initially treated with WT-BiTE were re-dosed with WT-, mut1-, or mut2-BiTE. Re-dosing with mut1-or mut2-BiTE significantly restored cytotoxicity against the residual tumor populations compared with re-dosing with WT-BiTE (**Figs. 5F–5G**). Together, these results demonstrate that affinity maturation expands the functional glycoform coverage of a single BiTE, enabling sustained elimination of heterogeneous Tn/STn tumor populations even as the composition shifts toward STn and thereby preventing the progressive loss of efficacy observed with the parental 5E5 BiTE.

To determine whether dual-glycoform recognition could improve antitumor activity in the context of glycoform heterogeneity, we evaluated mut1-BiTE in a PBMC-humanized mouse model bearing heterogeneous Tn/STn-expressing tumors. NCG mice were inoculated subcutaneously on day 0 with HT29 colon cancer cells (0.6 × 10^6^) composed of a 1:1 mixture of Tn-and STn-expressing populations, followed by engraftment with 6 × 10^6^ human PBMCs on day 1. Tumor growth was monitored by bioluminescence imaging beginning on day 3, after which mice were randomized into PBS, WT-BiTE, or mut1-BiTE treatment groups. Mice received 10 μg of BiTE on days 4 and 6, followed by 20 μg every other day beginning on day 8 (**Fig. 5H**). Mut1-BiTE induced robust and durable tumor suppression, as evidenced by significantly reduced bioluminescence signal and tumor volume throughout the study. By contrast, WT-BiTE failed to significantly control tumor growth relative to PBS, with both groups showing comparable tumor progression (**Figs. 5I**–**5M**). These results are consistent with the limited functional glycoform coverage of WT-BiTE observed in the heterogeneous in vitro models. Together, these findings demonstrate that enhanced dual MUC1-Tn/STn recognition by mut1-BiTE translates into improved antitumor efficacy *in vivo*.

### Affinity maturation enhances BiTE activity across heterogeneous patient-derived tumor-like cell clusters

To evaluate the translational potential of affinity-matured BiTEs, we employed patient-derived tumor-like cell clusters (PTCs)(53,54), the *ex vivo* tumor models which preserve the endogenous cellular architecture of primary tumors including tumor-infiltrating lymphocytes (TILs), enabling direct assessment of BiTE-mediated cytolytic activity within a clinically relevant patient-specific context without exogenous effector cell supplementation (**Fig. 6A**). We assessed mut1-BiTE and WT-BiTE against a panel of eighteen PTCs spanning multiple solid tumor specimens including lung, bladder, liver, pancreatic, gastric, intestinal, and colon cancers (**Fig. 6** and **Supplementary Figs S17-S19**). Prior to treatment, each PTC was characterized for target-antigen expression and endogenous immune-cell composition. MUC1-Tn and MUC1-STn expression was assessed by IHC staining (**Figs. 6L–6S**, and **Supplementary Fig S18**), while multiplex immunofluorescence staining for pan-cytokeratin (panCK; tumor cells), CD3 (T cells), and CD8 (cytotoxic T cells). T cell abundance was quantified based on CD8^+^/DAPI, CD3^+^/DAPI, and CD8^+^/CD3^+^ ratios across all specimens (**Supplementary Figs S17-S19** and **Table S2**).

**Figure 6.**
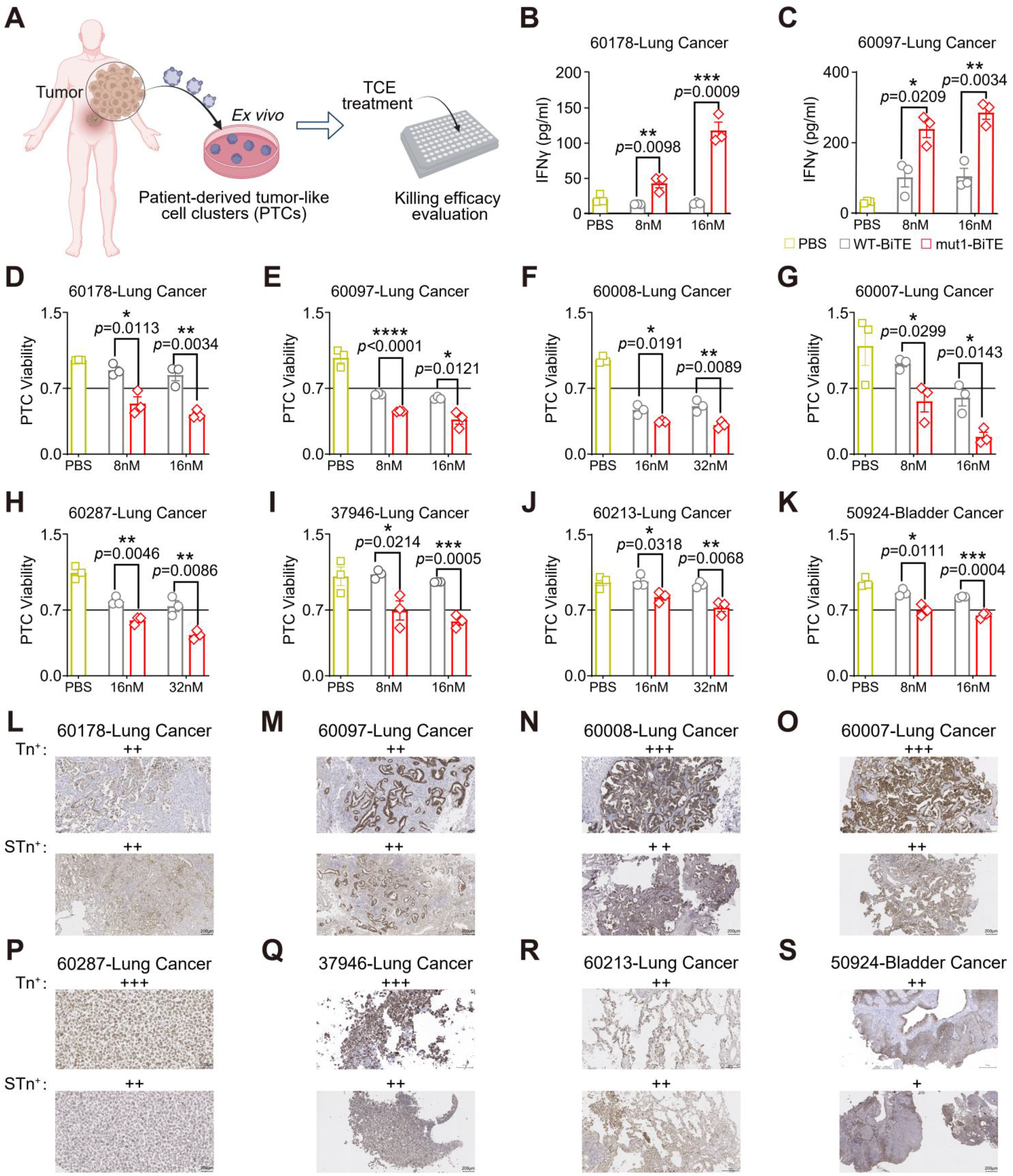
Affinity maturation enhances BiTE activity across heterogeneous patient-derived tumor-like cell clusters. **A)** Schematic illustration of the *ex vivo* patient-derived tumor-like clusters (PTC) platform used to evaluate the therapeutic efficacy of T-cell engagers (TCEs). Fresh tumor specimens were processed to establish PTCs, followed by treatment with the indicated BiTE molecules and assessment of antitumor activity. **B-C)** IFNγ secretion in representative PTC samples following treatment with PBS, WT-BiTE, or mut1-BiTE at the indicated concentrations. **D–K)** Viability of representative PTC samples from patients with lung cancer (60178, 60097, 60007, 60008, 37946, 60287 and 60213) and bladder cancer (50924) following treatment with PBS, WT-BiTE, or mut1-BiTE at the indicated concentrations. **L–S)** Representative immunohistochemical staining of Tn and STn in the corresponding patient tumor samples. Expression levels are indicated as negative (−), weak (+), moderate (++), or strong (+++) based on staining intensity and the proportion of positive cells. Data represent mean ± SEM; two-tailed unpaired Student’s *t*-test. ns, not significant; \**p* < 0.05; \*\**p* < 0.01; \*\*\**p* < 0.001; \*\*\*\**p <* 0.0001.

Eight of 18 PTCs demonstrated significant, concentration-dependent cytolytic responses to mut1-BiTE that were greater than those elicited by WT-BiTE (**Figs. 6D–6K**). These responding specimens, spanning seven lung cancer and one bladder cancer PTC, all exhibited moderate to strong MUC1-Tn and/or MUC1-STn expression by IHC (**Figs. 6L–6S**, Tn^++^ to ^+++^; STn^++^ to ^+++^) and harbored substantial endogenous cytotoxic TIL infiltration, quantified by CD3^+^/DAPI ratios ranging from 38% to 89% and CD8^+^/DAPI ratios ranging from 22% to 69% (**Supplementary Fig S17** and **Table S2**). Multiplex immunofluorescence revealed that CD3^+^ and CD8^+^ T cells were spatially co-localized with panCK^+^ tumor cell regions, indicating intratumoral T-cell infiltration rather than an exclusively peripheral or stromal distribution in all responding specimens (**Supplementary Fig S17**). Enhanced IFN-γ secretion in mut1-BiTE-treated PTCs relative to WT-BiTE further supported superior T cell activation in representative responding specimens (**Figs. 6B and 6C**). The magnitude of mut1-BiTE cytolytic activity was associated with MUC1-Tn/STn expression and the abundance of CD3^+^ and CD8^+^ TILs across individual PTCs, consistent with dependence on both target-antigen availability and endogenous effector-cell abundance.

In contrast, the remaining ten PTCs showed no cytolytic response to either mut1-BiTE or WT-BiTE. IHC and multiplex immunofluorescence revealed that non-response was associated with insufficient target-antigen expression and/or limited cytotoxic T-cell infiltration (**Supplementary Figs S18-S19**). The first group comprised bladder 50920 (Tn^−^/STn^+^), liver 60283 (Tn^+/−^/STn^+/−^), lung 60227 (Tn^+^/STn^+^), and pancreatic 19507 (Tn^+/−^/STn^+/−^) exhibited low or absent MUC1-Tn/STn expression together with relatively limited T-cell infiltration (CD3^+^/DAPI: 5–39%; CD8^+^/DAPI: 2.5–17%). The second nonresponsive group comprised specimens with moderate to strong MUC1-Tn/STn expression, including lung 60245 (Tn^+++^/STn^++^), intestinal 93830 (Tn^++^/STn^+++^), colon 02748 (Tn^+^/STn^++^), colon 02767 (Tn^+^/STn^++^), lung 60157 (Tn^++^/STn^++^), and gastric 02351 (Tn^++^/STn^++^), but exhibited markedly reduced cytotoxic T-cell infiltration, with CD8^+^/DAPI ratios of 0.08–7.1% compared with 22–69% in responding specimens. Notably, specimen 93830 exhibited substantial CD3^+^ T cell infiltration (CD3^+^/DAPI: 43.8%) but a markedly reduced CD8^+^/CD3^+^ ratio of only 8.5%, suggesting that T-cell composition, rather than the overall T-cell abundance, may influence BiTE responsiveness (**Supplementary Fig S19**). Collectively, these results indicate that mut1-BiTE responsiveness in patient-derived models is associated with the co-occurrence of sufficient MUC1-Tn/STn expression and CD8^+^ T-cell infiltration.

Together, these findings demonstrate that affinity maturation expands the functional activity of glycan-targeted BiTEs across heterogeneous patient-derived tumors while preserving target-antigen dependence, supporting dual MUC1-Tn/STn recognition as a strategy to address tumor glycoform heterogeneity.

## Discussion

Intratumoral heterogeneity remains a major obstacle to durable targeted cancer therapy because variation in antigen abundance or molecular identity can generate tumor-cell populations that are incompletely covered by a single therapeutic specificity(55,56). This challenge is especially relevant to tumor-associated glycans, whose dynamic biosynthesis produces closely related glycoforms that coexist within individual tumors(57,58). Here, we show that antibody affinity maturation can be used to address this form of heterogeneity. Directed evolution of 5E5 generated antibodies with high-affinity recognition of both MUC1-Tn and MUC1-STn while preserving glycoepitope specificity. We define the ability to maintain effective recognition and antitumor activity across coexisting glycoforms as functional glycoform coverage. Our findings reveal that affinity maturation can expand not only binding strength but also recognition breadth, providing a strategy to increase functional coverage across heterogeneous glycoform states.

The structural basis of this broadened recognition provides insight into how antibodies can accommodate closely related glycoforms without sacrificing epitope discrimination. Carbohydrate recognition is inherently challenging because glycans possess limited chemical diversity and often differ by subtle structural modifications(59,60). In the 5E5 system, relative preference for Tn versus STn does not arise primarily from extensive recognition of the additional sialic acid in STn. Instead, HCDR3 residue 98 fine-tunes glycoform preference through local, water-mediated interactions surrounding the glycosylated peptide. This arrangement allows a common binding architecture to accommodate both glycoforms while modulating their relative recognition. The recurrent contribution of LCDR1 Trp27d across distinct antibody scaffolds further suggests that strengthening interactions with the underlying peptide can provide a transferable strategy for enhancing MUC1 glycopeptide recognition.

The functional consequences of broadened recognition became most evident in heterogeneous tumor-cell populations. The affinity-matured BiTEs substantially reduced the concentration required for tumor-cell killing while producing more modest changes in maximal cytotoxicity, consistent with their slower dissociation rates and more sustained target engagement(61,62). More importantly, affinity maturation expanded the range of glycoform compositions over which productive T-cell killing could be maintained. Whereas WT-BiTE activity progressively deteriorated as tumors became increasingly STn dominant, mut1-and mut2-BiTEs retained robust activity across a broad range of Tn/STn ratios. Notably, increasing the proportion of poorly recognized STn-expressing cells reduced WT-BiTE-mediated killing not only of the STn ^+^ population but also of the remaining Tn ^+^ cells. Although the mechanism underlying this effect remains to be determined, these findings suggest that poorly engaged tumor-cell populations can compromise overall T-cell redirection within a heterogeneous tumor. The principal functional consequence of affinity maturation was therefore not simply increased potency, but expanded functional glycoform coverage.

This concept is related to, but mechanistically distinct from, emerging dual-targeting strategies designed to mitigate antigen heterogeneity and therapeutic escape. Dual-antigen therapeutics recognize two different tumor-associated targets to broaden tumor coverage or exploit target co-expression, as exemplified by the EGFR×HER3 bispecific ADC BL-B01D1 and the EGFR×cMet bispecific antibody amivantamab(63,64). Dual-epitope strategies similarly engage distinct epitopes on the same antigen to enhance avidity, receptor clustering, or coverage of heterogeneous antigen states. These approaches typically expand therapeutic breadth by incorporating two binding specificities. By contrast, our strategy broadens the recognition spectrum of a single paratope, enabling one antigen-binding site to accommodate closely related tumor-associated glycoforms. Functional glycoform coverage therefore represents a complementary approach to antigen heterogeneity that does not require a second antigen-or epitope-binding arm and may be particularly suited to structurally related antigen states generated by post-translational modification.

The patient-derived tumor-like cluster studies further place this recognition problem within the immune context of human tumors. Broader glycoform recognition was not sufficient to generate cytotoxicity in every specimen; responsiveness was associated with both MUC1-Tn/STn expression and the presence of CD8^+^ tumor-infiltrating lymphocytes. This is particularly relevant for T-cell engagers, whose activity requires both adequate tumor-cell recognition and access to functional effector cells(65). The observation that high total CD3^+^ infiltration did not consistently correspond with responsiveness further suggests that cytotoxic T-cell availability may be more relevant than overall T-cell abundance. Given the size and histologic diversity of the cohort, these findings should not yet be interpreted as defining predictive biomarkers. Rather, they point to two biologically distinct constraints—target availability and effector-cell availability—that may jointly determine whether enhanced molecular recognition can be translated into tumor killing. This framework may be useful for future patient-selection strategies in which both tumor glycoantigen expression and immune context are considered.

An important feature of the evolved antibodies is that broader glycoform recognition was achieved without detectable loss of tumor-associated specificity in the systems examined. This consideration is particularly important for T-cell engagers, for which recognition of low-level target expression in normal tissues can restrict the therapeutic window (66,67). Affinity maturation preserved dependence on the glycosylated MUC1 epitope and did not produce detectable binding across the normal tissues tested. The apical localization and more extensive O-glycosylation of MUC1 in normal glandular epithelia may provide additional layers of selectivity, although their contributions cannot be established from the present experiments(68,69). These findings suggest that recognition breadth and tumor-associated glycoepitope specificity need not be mutually exclusive. Nevertheless, broader normal-tissue profiling, evaluation of soluble or shed MUC1, and formal toxicology will be necessary to define the therapeutic window before clinical translation.

More broadly, our findings suggest that antibody engineering for heterogeneous tumor antigens may require balancing molecular specificity with recognition breadth. Conventional antibody optimization often seeks maximal discrimination of a single, precisely defined antigenic structure. Such an approach is advantageous when the target is uniform, but it can leave gaps in tumor coverage when closely related disease-associated molecular states coexist or interconvert. Dual-antigen and dual-epitope approaches address this problem by increasing the number of binding specificities within a therapeutic molecule. Our findings establish a complementary principle for structurally related antigen variants: a single binding site can be engineered to recognize a defined spectrum of disease-associated molecular states while preserving discrimination from normal antigen forms. For tumor-associated glycans, functional glycoform coverage provides a useful framework for considering this balance between specificity and breadth.

## Conclusion

In summary, affinity maturation of 5E5 expanded recognition across tumor-associated MUC1 glycoforms while preserving glycoepitope specificity. This broadened recognition translated into sustained T-cell redirection across heterogeneous tumor populations, including increasingly STn-dominant conditions under which the parental 5E5 BiTE progressively lost efficacy. By linking structural adaptation of a single paratope to functional activity across heterogeneous glycoform states, our findings establish functional glycoform coverage as an additional design parameter for glycan-directed cancer therapeutics. This concept complements conventional optimization of affinity and specificity and provides a framework for engineering therapies that are more resilient to intratumoral antigen heterogeneity.

## Data Availability

The data generated in this study are available within the article and its Supplementary Data. The atomic coordinates and structure factors for the antibody–ligand complexes 5E5 scFv/Ag_Tn, 5E5 mut2 Fab/Ag_Tn, 5E5 mut3 Fab/Ag_Tn, and 5E5 mut3 Fab/Ag_STn have been deposited in the Protein Data Bank (PDB) under accession codes 38SP, 38SQ, 38SR, and 38SS, respectively. Additional data supporting the findings of this study are available from the corresponding author upon reasonable request.

## Authors’ contribution

J.Y. and P.W. conceived the study and designed the research with J.J. J.Y. performed the majority of the experiments, analyzed and interpreted the data, and prepared the manuscript. X.Z. and I.A.W. contributed to the crystallographic studies and structural analyses. H.L. and J.J.X. contributed to the patient-derived tumor-like cell cluster (PTC) experiments. J.C. and J.J. contributed to the construction of the antibody libraries. D.Z. performed the biolayer interferometry experiments. Y.T. and L.W. contributed to the synthesis of glycopeptides. Z.Y., S.C., and L.-A.C. assisted with colony construction. J.Z. assisted with tissue processing. P.W. supervised the overall study and provided funding and resources. J.Y. and P.W. wrote the manuscript with input from all authors. All authors reviewed and approved the final manuscript.

## Acknowledgements

We thank Henry Tien for automated robotic crystal screening. X-ray diffraction datasets were collected at the Stanford Synchrotron Radiation Lightsource (SSRL) beamlines 12-1. Use of the Stanford Synchrotron Radiation Lightsource, SLAC National Accelerator Laboratory, is supported by the U.S. Department of Energy, Office of Science, Office of Basic Energy Sciences under Contract No. DE-AC02-76SF00515. The SSRL Structural Molecular Biology Program is supported by the DOE Office of Biological and Environmental Research, and by the National Institutes of Health, National Institute of General Medical Sciences (P30GM133894). The contents of this publication are solely the responsibility of the authors and do not necessarily represent the official views of NIGMS or NIH. Peng Wu, Joseph Jardine and Ian Wilson are partially supported by Scripps Research.

