## Supplementary Information for "Affinity Maturation Enables Dual Targeting of Cancer-Associated MUC1-Tn/STn and Drives Potent Tumor Control"

for

### Key resources table

### Materials and Methods

#### Supplementary Figures S1 to S20

**Figure S1.** Overall structure of 5E5 scFv in complex with MUC1-Tn glycopeptide Ag\_Tn (GST(Tn)APPA)

**Figure S2.** Flow cytometry gating strategy for yeast display selection

**Figure S3.** Yeast display screening and enrichment of antibody variants against MUC1-Tn and MUC1-STn glycoforms

**Figure S4.** Enhanced recognition of MUC1-Tn and MUC1-STn peptides by affinity-matured 5E5 variants

**Figure S5.** Epitope specificity profiling and tissue cross-reactivity analysis of WT and affinity-matured 5E5 variants

**Figure S6.** Electron density maps for peptides Ag\_Tn or Ag\_STn in their complexes with 5E5\_mut2 or 5E5\_mut3 Fab

**Figure S7.** Mutated residues of the affinity-matured 5E5-mut2 and 5E5-mut3 Fabs as observed in their complex structures with Ag\_Tn

**Figure S8.** Structural comparison of 5E5-mut2 or 5E5-mut3 Fabs with 5E5 WT scFv in their complex with Ag\_Tn around mutated HCDR residues 33 and 51

**Figure S9.** Structural comparison of 5E5-mut2 or 5E5-mut3 Fabs with 5E5 WT scFv around mutated H-chain residue 102 as observed in their complex with Ag\_Tn or Ag\_STn

**Figure S10.** Structural basis of 5E5\_mut2 or 5E5\_mut3 Fabs with 5E5 WT scFv around mutated H-chain residue 98 as observed in their complex with Ag\_Tn or Ag\_STn

**Figure S11.** Structural basis of 5E5\_mut2 and 5E5\_mut3 Fabs with 5E5 WT scFv around L-chain residues 50 and 52 as observed in their complex with Ag\_Tn or Ag\_STn

**Figure S12.** Affinity-enhancing N/Y<sup>27d</sup>W mutations do not introduce non-specific binding to normal human cells across multiple anti-MUC1 antibody scaffolds.

**Figure S13.** BiTE linker optimization and T cell activation across MUC1-Tn/STn-expressing cancer cell lines

**Figure S14.** Mut1-BiTE enhances T cell cytokine production and cytotoxic effector function across MUC1-Tn/STn-expressing cancer cell lines

**Figure S15.** Bispecific T cell engagers derived from affinity-matured 5E5 variants display enhanced cytolytic activity

**Figure S16.** Affinity-matured MUC1 BiTEs enhance T-cell degranulation across different Tn/STn antigen ratios

**Figure S17.** Immunofluorescence characterization of T-cell infiltration in BiTE-responsive patient-derived tumor samples

**Figure S18.** Representative non-responsive PTC samples to WT and affinity-matured MUC1 BiTEs

**Figure 19.** Immunofluorescence characterization of immune-cell infiltration for nonresponsive PTCs

**Figure S20.** SDS-PAGE analysis confirming purity of affinity-matured 5E5 Fc-fusion, Fab, and BiTE proteins

#### Supplementary Table S1: X-ray data collection and refinement statistics

#### Supplementary Table S2: Quantification of tumor-infiltrating lymphocyte composition across eighteen PTC specimens

### References

### Key resources table

| REAGENT or RESOURCE | SOURCE | IDENTIFIER |
| --- | --- | --- |
| **Antibodies** |  |  |
| Anti-human CD3 (HIT3a) | BioLegend | 300306 |
| Anti-human CD8 (HIT8a) | BioLegend | 300912 |
| Anti-human CD4 (RPA-T4) | BioLegend | 300521 |
| Anti-human CD107a (H4A3) | BioLegend | 328608 |
| Anti-human Granzyme B (QA16A02) | BioLegend | 372228 |
| Anti-human CD25 (M-A251) | BioLegend | 356104 |
| Anti-human EpCAM (W21090A) | BioLegend | 624152 |
| Anti-human MUC1 (DF3) | BioLegend | 311203 |
| Anti-human IgG Fc (QA19A42) | BioLegend | 366914 |
| Anti-rat IgG Fc (Poly4054) | BioLegend | 405420 |
| Streptavidin | BioLegend | 405237 |
| Anti-human IgG Fc (RRID: AB_2337882) | Jackson ImmunoResearch | 109-605-008 |
| Anti-Mouse IgG Fc (RRID: AB_2338842) | Jackson ImmunoResearch | 115-545-008 |
| Anti-Biotin (RRID: AB_2339046) | Jackson ImmunoResearch | 200-602-211 |
| Anti-c-Myc antibody (polyclonal) | Abcam | ab1394 |
| Anti-V5 Epitope Tag antibody (1036H) | R&D Systems | FAB8926V |
| Anti-human IgG-HRP | Thermo Fisher | A18817 |
| Anti-human Fab | Fisher Scientific | NC0201453 |
| **Chemicals, peptides, and recombinant proteins** |  |  |
| Vicia Villosa Lectin (VVL, VVA), Biotinylated | Vector Laboratories | B-1235-2 |
| D-Luciferin | GoldBio | eLUCNA-2G |
| Protein A resin | VWR | 76510-282 |
| HisPur™ Ni-NTA Resin | Life Technologies | 88221 |
| EZ-Link™ NHS-Biotin | Thermo Fisher | 20217 |
| Cas-9 nuclease | IDT | 1081058 |
| Gene synthesis | Twist Bioscience | N/A |
| Gene Oligo pool synthesis | IDT | N/A |
| Bacto™ Casamino Acids | Gibco | 223050 |
| D-Galactose | Sigma | G0625 |
| D-Glucose | Sigma | G7021 |
| YPD Broth | BD Biosciences | 242820 |
| D-Sorbitol | Thermo Scientific | 036404.36 |
| SD-Ura | Sunrise Science Products | 1703-500 |
| 0.1M Glycine beffer | Teknova | G4527 |
| Yeast Nitrogen Base | Sigma | Y0626 |
| L-Tryptophan | MilliporeSigma | 6540 |
| Na<sub>2</sub>HPO<sub>4</sub>·7H<sub>2</sub>O | MilliporeSigma | S9390 |
| NaH<sub>2</sub>PO<sub>4</sub>·2H<sub>2</sub>O | MilliporeSigma | 04269 |
| Sodium citrate | Sigma | S1804 |
| **Experimental models: Cell lines** |  |  |
| Jurkat | This paper | N/A |
| HEK293T | This paper | N/A |
| OVCAR3 | This paper | N/A |
| MCF7 | This paper | N/A |
| T-47D | This paper | N/A |
| HT29 (COSMC KO) | This paper | N/A |
| MDA-MB-231 (COSMC KO) | This paper | N/A |
| HT29 (ST6GalNAc OE) | This paper | N/A |
| MDA-MB-231 (ST6GalNAc OE) | This paper | N/A |
| Expi293 | This paper | N/A |
| *Saccharomyces cerevisiae* YVH10 | Joseph Jardine Lab | N/A |
| 10-beta Competent *E. coli* | NEW ENGLAND Biolabs | C3019H |
| **Experimental models: Organisms/strains** |  |  |
| Female NCG mice | Charles River | N/A |
| **Critical commercial assays** |  |  |
| ELISA MAX™ Deluxe Set Human IL-2 | BioLegend | 431804 |
| ELISA MAX™ Deluxe Set Human IFN-γ | BioLegend | 430104 |
| BCA Protein Assay Kit | Thermo Fisher | 23227 |
| Frozen-EZ Yeast Transformation II Kit | Zymo Research | T2001 |
| Zymoprep Yeast Plasmid Miniprep II | Zymo Research | D2004 |
| Monarch® Spin Plasmid Miniprep Kit | NEW ENGLAND Biolabs | T1110L |
| Midi kit for endotoxin-free plasmid DNA | Fisher Scientific | NC2220631 |
| ExpiFectamine 293 transfection reagents | Life Technologies | A14524 |
| **REAGENT or RESOURCE** |  |  |
| Lipofectamine™ 2000 | Thermo Fisher | 11668027 |
| Tween 20 | Sigma | P1379 |
| Bovine Serum Albumin | Fisher Scientific | BP1600100 |
| DMEM | Thermo Fisher | 10569044 |
| RPMI1640 | Life Technologies | 21870092 |
| HEPES | Gibco | 15630-080 |
| Glutamax | Gibco | 35050061 |
| Penicillin-Streptomycin | Gibco | 15140122 |
| Sodium Pyruvate | Gibco | 11360070 |
| Non-Essential Amino Acids | Gibco | 11140050 |
| 2-Mercaptoethanol | Life Technologies | 21985023 |
| Fetal Bovine Serum | Phoenix Scientific | PS-100 |
| Expi293 expression medium | Thermo Fisher | A1435102 |
| HiFi DNA Assembly Master Mix | NEB | E2621S |
| Normal Goat Serum | Jackson Immuno Research | 005-000-121 |
| Human Breast Cancer Tissue | Tissuearray | HuCAT297 |
| Human Colon Cancer Tissue | Tissuearray | HuCAT126 |
| Normal Human Skin Tissue | Tissuearray | HuFPT136 |
| Human Multi-normal organs tissue array | Tissuearray | FDA331a |
| Multiple organ of tumor and normal tissue array | Tissuearray | CTRL301a |
| FUW-Luc-mCherry-puro | Addgene | 183685 |
| TMB Solution (1X) | Life Technologies | 00-4201-56 |
| BsaI-HF®v2 | NEW ENGLAND Biolabs | R3733S |
| BsmBI-v2 | NEW ENGLAND Biolabs | R0739S |
| EcoRI-HF® | NEW ENGLAND Biolabs | R3101S |
| NheI-HF® | NEW ENGLAND Biolabs | R3131S |
| KLD Enzyme Mix | NEW ENGLAND Biolabs | M0554S |
| DPBS, no calcium, no magnesium | Gibco | 14190250 |
| Trypsin-EDTA (0.25%) | Gibco | 25200114 |
| **Software and algorithms** |  |  |
| FlowJoX | FlowJo, LLC | N/A |
| GraphPad Prism | GraphPad | N/A |
| IVIS Living Image |  | N/A |
| Biorender | biorender.com | N/A |
| Adobe Illustration | Adobe Creative Cloud | N/A |
| IncuCyte S3 | Sartorius | N/A |
| Synergy H4 Hybrid Reader | BioTek | N/A |
| PyMol | Schrödinger, LLC | N/A |
| Phaser | McCoy et al. <sup>1</sup> | PMID 15805601 |
| Coot | Emsley et al. <sup>2</sup> | PMID 20383002 |
| Refmac5 | Murshudov et al. <sup>3</sup> | PMID 15299926 |
| Phenix | Adams et al. <sup>4</sup> | PMID 20124702 |
| MolProbity | Chen et al. <sup>5</sup> | PMID 20057044 |
| HKL2000 | Otwinowski and Minor <sup>6</sup> | PMID 27754618 |
| **Deposited data** |  |  |
| 5E5 WT_scFv in complex with Ag_Tn | This paper | PDB: 38SP |
| 5E5 mut2_Fab in complex with Ag_Tn | This paper | PDB: 38SQ |
| 5E5 mut3_Fab in complex with Ag_Tn | This paper | PDB: 38SR |
| 5E5 mut3_Fab in complex with Ag_STn | This paper | PDB: 38SS |

### Materials and Methods

#### Human samples

Human tumor specimens used for patient-derived tumor-like cell cluster (PTC) studies were obtained from Beijing Cancer Hospital and Peking University People's Hospital. This study was approved by the medical ethics committee of Peking University Cancer Hospital (No. 2019KT41, No. NCT05280210), Peking University People's Hospital (No. 2021PHB239-001). Before surgery, all patients provided written informed consent for the use of excess tissue for research purposes. The collection and use of these specimens were approved by the relevant institutional ethics committees in China. Whole blood from healthy donors used for the isolation of primary human T cells was obtained from the Normal Blood Donor Service (NBDS) at Scripps Research under an Institutional Review Board (IRB)-approved protocol.

#### Animals

Female NCG (NOD-*Prkdc*<sup>em26Cd52</sup>//*2rg*<sup>em26Cd22</sup>/NjuCrl, strain: 572) mice were obtained from the Charles River Laboratories. Seven-week-old female mice were used for this study. Mice were housed in ventilated, sterile-barrier cages under pathogen-free conditions in the Immunology Vivarium of the Department of Animal Resources at Scripps Research. All animal experiments were conducted under protocols approved by the Institutional Animal Care and Use Committee (IACUC) of The Scripps Research Institute.

#### Cell lines

All cell lines were maintained under standard conditions in a humidified incubator at 37°C with 5% CO<sub>2</sub>. All human cancer cell lines were cultured in complete medium supplemented with 10% FBS (Phenix), 1% Penicillin-Streptomycin, 1% GlutaMAX, 1% HEPES, 1% non-essential amino acids, and 1% sodium pyruvate. OVCAR3 and Jurkat cells were cultured in complete RPMI-1640 medium. MCF7, T-47D, HT29, MDA-MB-231, and HEK293T cells were grown in complete DMEM. Human PBMCs were cultured in complete RPMI-1640 medium supplemented with 0.01% 2-mercaptoethanol. Expi293 cells were maintained in Expi293™ Expression Medium at 37°C in a humidified shaker at 8% CO<sub>2</sub> and 150 rpm. *Saccharomyces cerevisiae* YVH10 was cultured in YPD medium at 30°C with shaking at 225 rpm.

#### Crystal structure determination.

All crystallization experiments of 5E5 WT scFv, mut2 and mut3 Fabs in complex with Ag\_Tn or Ag\_STn were set up using the sitting-drop vapor diffusion method with automated crystal screening on our robotic Rigaku CrystalMation system at Scripps Research. For each complex, 0.1 μL of protein pre-incubated with the respective antigen at a molar ratio of 1:5 was mixed with 0.1 μL of well solution at 20°C, and crystals were flash-cooled at 100 K prior to data collection. Specific conditions were as follows: the 5E5 scFv/Ag\_Tn complex (7.0 mg/ml protein in 20 mM Tris pH 8.0, 150 mM NaCl, and 0.02% (v/v) NaN<sub>3</sub>; well solution: 0.1 M Tris pH 8.67 and 2.4 M ammonium sulfate; cryoprotectant: 20% (v/v) ethylene glycol); the 5E5-mut2 Fab/Ag\_Tn complex (10.0 mg/ml protein in the same buffer; well solution: 0.1 M sodium citrate pH 5.6, 0.17 M ammonium sulfate, 15% (v/v) glycerol, and 25.5% (w/v) polyethylene glycol 4,000; no additional cryoprotectant); the 5E5-mut3 Fab/Ag\_Tn complex (10.0 mg/ml protein in the same buffer; well solution: 0.2 M lithium citrate and 20% (w/v) polyethylene glycol 3,350; cryoprotectant: 20% (v/v) ethylene glycol); and the 5E5-mut3 Fab/Ag\_STn complex (7.1 mg/ml protein in the same buffer; well solution: 0.1 M sodium citrate pH 5.6, 0.2 M ammonium acetate, and 30% (w/v) polyethylene glycol 4,000; cryoprotectant: 10% (v/v) ethylene glycol). Diffraction data were collected at the Stanford Synchrotron Radiation Lightsource (SSRL) beamline 12-1 (Table S1). Data for all crystals were integrated and scaled with HKL2000<sup>6</sup>. Data collection statistics are summarized in Table S1.

Crystal structures of all the four complexes were determined by molecular replacement (MR) using the program Phaser<sup>1</sup> using the wild-type 5E5 scFv structure (PDB code 6TNP) as the input MR model. Initial rigid body refinement was performed in REFMAC5<sup>3</sup>, and subsequently simulated annealing and restrained refinement (including TLS refinement) were carried out in Phenix<sup>4</sup>. Between rounds of refinements, model building was examined and modified with the program Coot<sup>2</sup>. Final statistics for these structures are summarized in Table S1. The quality of the structures was analyzed using the JCSG validation suite (qc-check.usc.edu/QC/qc_check.pl) and MolProbity<sup>5</sup>. All figures were generated with PyMol (www.pymol.org).

### Yeast transformation

A single colony of *Saccharomyces cerevisiae* YVH10 cells (provided by the Jardine laboratory, Scripps Research) was inoculated into 2 mL YPD medium (cat. no. 242820, filter-sterilized) and grown overnight at 30°C with shaking. The overnight culture was diluted into 50 mL fresh YPD medium and grown at 30°C with shaking until OD<sub>600</sub> reached approximately 1.5. Cells were harvested by centrifugation and resuspended in 25 mL of 100 mM lithium acetate. Dithiothreitol (DTT; 250 µL of a 1 M stock solution) was added, mixed rapidly, and incubated at 30°C for 10 min with shaking. Cells were then pelleted, washed once with 25 mL of ice-cold deionized water, and resuspended in ice-cold deionized water to a final volume of 500 µL. The cell suspension was divided equally into two microcentrifuge tubes (250 µL each). To each tube, 1 µg of linearized vector DNA and 5 µg of insert library DNA were added and mixed by gentle pipetting. Each cell–DNA mixture was transferred to a 0.2 cm gap electroporation cuvette (Bio-Rad) and electroporated using a Gene Pulser Xcell system (Bio-Rad) with the following parameters: square wave mode; 500 V; pulse length 15.0 ms; single pulse; 2 mm cuvette. Immediately after electroporation, cells were transferred to 25 mL YPD medium and allowed to recover at 30°C for 1 h with shaking at 200 rpm. To estimate transformation efficiency, 2.5 µL of the recovery culture was removed and plated as a dilution series for colony-forming unit (CFU) enumeration. The remaining culture was transferred to 1 L of SD-Ura selective medium (without tryptophan; Sunrise Science) supplemented with 1% Penicillin-Streptomycin and grown overnight at 30°C with shaking.

### Yeast library labeling and sorting

Following transformation, yeast cells were expanded and passaged once to improve surface display efficiency. For induction, cells were grown overnight at 30°C in SGCAA induction medium (1 L deionized water containing 20 g galactose, 1 g glucose, 6.7 g yeast nitrogen base without amino acids, 5 g bacto casamino acids, 10.1 g Na<sub>2</sub>HPO<sub>4</sub>·7H<sub>2</sub>O, 11.18 g NaH<sub>2</sub>PO<sub>4</sub>·2H<sub>2</sub>O, and 85.6 mg L-tryptophan; pH 6.5; filter-sterilized). For each library, 5×10<sup>7</sup> yeast cells were stained per sample in the first round of selection; 1×10<sup>7</sup> cells were used in all subsequent rounds. Cells were pelleted by centrifugation, washed with PBS containing 1% BSA (PBS/BSA), and incubated with biotinylated MUC1-Tn or MUC1-STn glycopeptide antigen at several non-depleting concentrations for at least 30 min at 4°C. After washing, cells were stained with FITC-conjugated anti-c-Myc antibody (Abcam, cat. no. ab1394), Alexa Fluor 405-conjugated anti-V5 antibody (R&D Systems, cat. no. FAB8926V), and Alexa Fluor 647-conjugated anti-biotin antibody (Jackson ImmunoResearch, cat. no. 200-602-211), each at 1:100 dilution, for 20 min at 4°C. After washing, cells were resuspended in 1 mL PBS/BSA and performed sorting. The top 5–10% of cells with the highest binding signal at a defined antigen concentration were collected. Sorted cells were expanded overnight at 30°C in 2 mL SD-Ura selective medium supplemented with 1% Penicillin-Streptomycin.

### Expression and Purification of Fab, scFv, scFv-Fc and BiTE proteins

Genes encoding the Fab, BiTE, scFv and scFv-Fc constructs were synthesized in IDT/Twist Bioscience and subcloned into the pFUSE expression vector. Proteins were produced by transient transfection of Expi293 cells. Fab, scFv and BiTE proteins were purified using a His-tag (6×His) affinity column, and scFv-Fc proteins were purified using a Protein A affinity column. Protein-containing fractions were pooled, concentrated, and buffer-exchanged into PBS using 0.5 mL Amicon Ultra centrifugal filters with a 10 kDa molecular weight cutoff (Merck Millipore). Protein purity was assessed by reducing or non-reducing SDS-PAGE. Protein concentrations were determined by BCA assay (Thermo Fisher). Purified proteins were aliquoted and stored at –80°C for long-term storage or –20°C for short-term use.

### PBMC Isolation

PBMCs were isolated from whole blood obtained from healthy donors through the Normal Blood Donor Service (NBDS) of The Scripps Research Institute. Briefly, blood samples were diluted 1:1 with DPBS containing 2 mM EDTA and 1% FBS, and carefully layered onto Ficoll-Paque Plus (GE Healthcare, cat. no. 17-1440-02) for density gradient separation. Samples were centrifuged at 1,200 × g for 20 min with minimum acceleration and deceleration. The mononuclear cell layer (buffy coat) was collected and washed twice with DPBS supplemented with 2 mM EDTA and 1% FBS.

### ELISA Binding assay

96-well high-binding microplates were coated with 200 ng MUC1 peptides or glycosylated MUC1 peptide in DPBS buffer at 4 °C overnight. After washing and blocking, plates were incubated with serial dilutions of 5E5 WT-Fc and mutant scFv-Fc fusion proteins at r.t. for 2 h, followed by incubation with goat anti-human IgG Fc-HRP secondary antibody at r.t. for 1 h. TMB colorimetric substrate was added to each well and incubated at r.t. for 15 min. The reaction was stopped by addition of 2 N H<sub>2</sub>SO<sub>4</sub>. Absorbance was measured at 450 nm using a plate reader.

#### **Biolayer Interferometry (BLI) assay**

BLI measurements were performed on an Octet RED96 instrument (ForteBio) using streptavidin (SA) biosensors (Sartorius, #18-5019). All samples were reconstituted in kinetics buffer (PBS containing 0.05% Tween-20). Biotinylated Ag\_Tn and Ag\_STn antigens were loaded onto SA biosensors at 10 µg/ml for 3 min. Association was measured by immersing the loaded biosensors into twofold serial dilution of Fab (5E5 WT\_Fab start at 1 µM; mut1-4 start at 250 nM) for 3 min, followed by dissociation in kinetics buffer for 5 min. All signals were reference-subtracted, and kinetic parameters ($K_D$, $k_{on}$ and $k_{off}$) were determined by global fitting using the Octet Data Analysis software.

#### **Generation of HT29-Tn and MDA-MB-231-Tn Cell Lines**

COSMC knockout in wild-type HT29 and MDA-MB-231 cells was performed by CRISPR/Cas9 gene editing. Surface expression of the Tn antigen was confirmed by VVA lectin staining. The resulting COSMC-knockout cell lines are referred to throughout as HT29-Tn and MDA-MB-231-Tn.

#### **Generation of HT29-STn and MDA-MB-231-STn Cell Lines**

To generate stable cell lines overexpressing STn glycans, the *ST6GalNAc1* coding sequence (Accession#: Q9NSC7) was subcloned into the pFUW lentiviral expression vector. Lentiviral particles were produced by Lipofectamine™ 2000-based co-transfection of HEK293T cells with pFUW-ST6GalNAc1, pMD2.G, pRSV-Rev, and pMDLg/p. For transduction, HT29-Tn and MDA-MB-231-Tn cells were seeded in complete medium in 50 mm<sup>2</sup> dishes. At 70–80% confluency, the medium was replaced with fresh complete medium containing 20 µL of *ST6GalNAc1* lentiviral supernatant and 8 µg/mL polybrene (Sigma-Aldrich). After two passages, stably transduced cells were stained with the anti-STn antibody CC49 followed by fluorescence-activated cell sorting (FACS). The resulting stable cell lines are referred to throughout as HT29-STn and MDA-MB-231-STn.

#### **Flow cytometry**

Anti-human antibodies to CD3 (HIT3a), CD4 (RPA-T4), CD8a (HIT8a), CD25 (M-A251), MUC1 (DF3), CD107a (H4A3), Granzyme B (QA16A02) and human IgG Fc (QA19A42) were purchased from BioLegend. Anti-biotin antibody (RRID: AB\_2339046) was from Jackson ImmunoResearch. Anti-c-Myc antibody (polyclonal) was from Abcam, Anti-V5 Epitope Tag antibody (1036H) was from R&D Systems. Anti-human Fab antibody (Alexa Fluor 647 AffiniPure Goat Anti-Human IgG, F(ab')<sub>2</sub> fragment specific) was from Fisher Scientific. VVA lectin was from Vector Laboratories. For cell surface binding experiments, Fab, BiTE, or scFv-Fc proteins were pre-mixed with either AF647-conjugated anti-human Fab or AF647-conjugated anti-human IgG Fc secondary antibody and incubated with cells on ice for 30 min in the dark. For T cell activation experiments, anti-human CD3 and CD25 antibodies incubated with cells on ice for 30 min in the dark. For CD107a and Granzyme B measurement, refer the protocol described as<sup>1, 2</sup>, Flow cytometry data were acquired on an Attune NxT Acoustic Focusing Cytometer (Thermo Fisher Scientific).

#### **In Vitro Cytotoxicity Assay**

Target tumor cells were seeded into 96-well flat-bottom plates at 10,000 cells per well. After allowing cells to adhere for 3 h, PBMCs and BiTE constructs at the indicated concentrations were added at an effector-to-target ratio of 5:1 to 8:1. Plates were immediately transferred to an Incucyte® Live-Cell Analysis System and imaged at 3 h intervals. T cell-mediated cytotoxicity was quantified by normalizing GFP or RFP fluorescence intensity to the PBS control at each time point.

#### **In vivo experiments**

NCG mice were inoculated subcutaneously (s.c.) with a 1:1 mixture of HT29-Tn and HT29-STn cells. Following tumor establishment, mice were engrafted intravenously (i.v.) with $8 \times 10^6$ primary human PBMCs. Three days after PBMC engraftment, mice were randomly grouped and treated with PBS (vehicle control) or BiTEs construct via i.v. injection every two days. Tumor burden was monitored by bioluminescence imaging using Living Image software and quantified by caliper measurement of tumor volume.

#### **Immunofluorescence (IF) staining of commercial human tissues**

Formalin-fixed, paraffin-embedded (FFPE) human tissue sections were deparaffinized in xylene (three washes) and rehydrated through a graded ethanol series. Antigen retrieval was performed by heating sections in 20 mM citrate buffer (pH 6.0). Sections were blocked with 5% BSA and 5% normal goat serum in PBST at room temperature for 1 h. Sections were then incubated overnight at 4°C with primary anti-MUC1 antibody (clone DF3) together with 5E5 scFv-Fc or mut1 scFv-Fc fusion protein. After washing, sections were incubated with fluorescent secondary antibodies at room temperature for 1 h, followed by nuclear counterstaining with DAPI. Images were acquired by fluorescence microscope.

#### **Establishment of the PTC models**

The PTC models were established as described previously. Collected fresh patient samples were conditioned in ice-cold PBS with 10 mM HEPES and 100 U/mL penicillin-streptomycin. Necrotic areas and adipose tissue were removed as possible. Tissues were minced and digested into small pieces in 5 ml collecting buffer containing 1 mM EDTA and 200 U/mL collagenase I, II and IV (Thermo Fisher Scientific) in PBS at 37 °C for 1 hour. 40 µm filters were used to collect dissociated cells. The suspension cells were centrifuged at 300 g, 4°C for 10 mins, then cell pellets were re-suspended in PTC growth medium. Malignant pleural effusions and ascites were centrifuged at 300 g for 5 min, and cell pellets were resuspended in PBS and added over Lymphocyte Separation Medium (MPbio) in centrifuge tube. After centrifugation at 400 g for 20 mins, middle cell layer was collected and resuspended in PTC growth medium. All cell suspensions from surgical resections, ascites or pleural effusions in PTC growth medium were seeded in low-attachment-surface dish at the concentration of $10^5$ cells/cm<sup>2</sup> and cultured in incubator at 37°C, 5% CO<sub>2</sub>. PTC growth medium was refreshed every 2-3 days.

#### **PTC viability detection**

The PTC viability assay was conducted in 96-well cell culture plates following the procedure described previously. PTCs more than 40 µm in diameter were collected using 40-µm filters and resuspended in PTC growth medium. 100 µL of a medium containing 30-50 PTCs was seeded into Teflon-modified chip (GeneX Health, Cat# GX-01). Next, 10 nM CD3-PDL1 BiTE, 8nM, 16 nM, 32 nM 5E5 WT or mut1 BiTE or the same volume of PBS were added to the well of PTCs. PTCs were incubated at 37 °C, 5% CO<sub>2</sub> for 7 days. All clusters were photographed at day 0 and day 7 in each well using the Nikon Ti-U microscope system. The drug effect by measuring the area of all PTCs in a whole well. The cell viability after applying BiTEs or PBS treatment was calculated according to the following formula:

$$p(x) = \frac{S_{t1}}{S_{t0}}, \quad p = \left(\frac{1}{n}\right) \sum_{i=1}^n p(x)$$

where S is the sum of cluster areas in a well and t0 and t1 are the time points when the areas are measured. p(x) is the PTC cell viability in each well, n is the number of the replicates, p is the average PTC cell viability of all replicates. The same cell viability value of the negative control (PBS) as a quality control. If the cell viability score p<sub>NC</sub> was less than 0.9, indicating that the PTC was possibly in the decline phase, the PTC test was discarded. The PTC areas were analyzed and quantified by the software ImageJ.

#### **Immunohistochemistry (IHC) staining of PTC samples**

PTCs were collected by centrifugation at 300g and 4 °C for 10 minutes. Tissues and PTCs were washed with cold PBS, and fixed in PBS containing 4% paraformaldehyde overnight. The pellets were paraffin-embedded, and 5 µm thick paraffin sections were generated and were dewaxed and rehydrated through a graded ethanol series. After heat-mediated antigen retrieval with Citrate Unmasking Solution (Solarbio life sciences, C1032), the sections were blocked at room temperature for 1 hour. The primary antibodies 5E5 scFv-Fc or CC49 scFv-Fc fusion protein were diluted in 3%

BSA, and staining was performed overnight at 4 °C with gentle rocking. DAB substrate kit (ZSGB-BIO, ZLI-9019) was used to provide an acceptable staining intensity. The stained tissue was visualized using the TEKSQRAY Slide Scanning System SQS-40P.

#### **IF staining of PTC samples**

The generated 5 µm thick paraffin sections of PTCs were dewaxed and rehydrated through a graded ethanol series. After heat-mediated antigen retrieval with Citrate Unmasking Solution, PTCs were permeabilized with 1% Triton-X 100 in PBS for 30 minutes, and blocked with 10% goat serum for 30 minutes. Sections were then incubated with primary antibodies (CD3, Abcam, ab11089; CD8a, Thermo Fisher Scientific, MA5-14548; panCK, Thermo Fisher Scientific, 53-9003-80) at 4 °C overnight followed by staining with Alexa 594 (Abcam, ab150160) or Alexa 647 (Abcam, ab150079)-conjugated secondary antibody. Nuclei were stained with DAPI (YEASEN, 40728ES03). Images were obtained with ZEISS AxioScan 7. The software ImageJ was used for image analysis.

#### **Statistics**

All figures are representative of at least three independent experiments unless otherwise noted. All graphs report mean ± SEM of biological replicates. Statistical significance of two-group comparisons was calculated using Student's t-test.

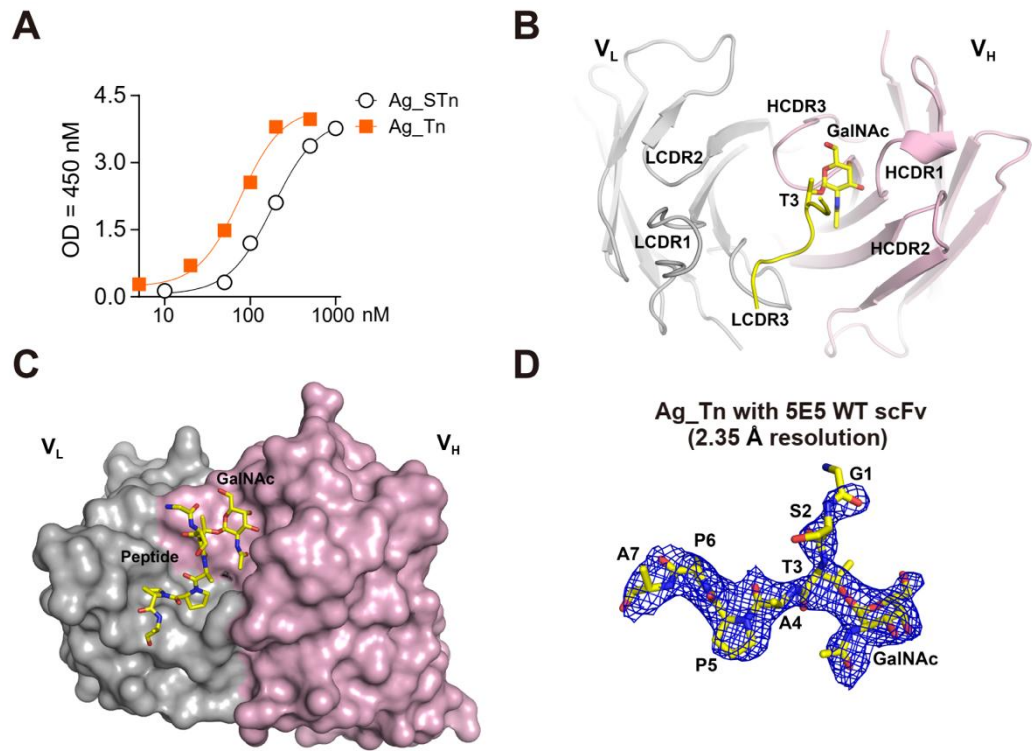

**Figure S1. Overall structure of 5E5 scFv in complex with MUC1-Tn glycopeptide Ag\_Tn (GST(Tn)APPA).**

**A**) ELISA binding curves of parental 5E5 scFc-Fc against MUC1-Tn (Ag\_Tn, orange squares) and MUC1-STn (Ag\_STn, gray circles) glycopeptides

**B**) Crystal structure of 5E5 scFv (grey: light chain; and light pink: heavy chain) bound to glycopeptide Ag\_Tn (yellow: carbon; red: oxygen; blue: nitrogen) with the Thr3 residue and GalNAc moiety in sticks.

**C**) Surface presentation of 5E5 scFv with Ag\_Tn displayed as sticks, demonstrating the paratope-epitope interface and antigen accommodation within the binding site.

**D**) Unbiased 2Fo-Fc electron density omit map (blue mesh, contoured at $1\sigma$) overlaid with the refined structure of Ag\_Tn shown as sticks, validating the quality of the model and confirming accurate positioning of the glycopeptide within the binding site.

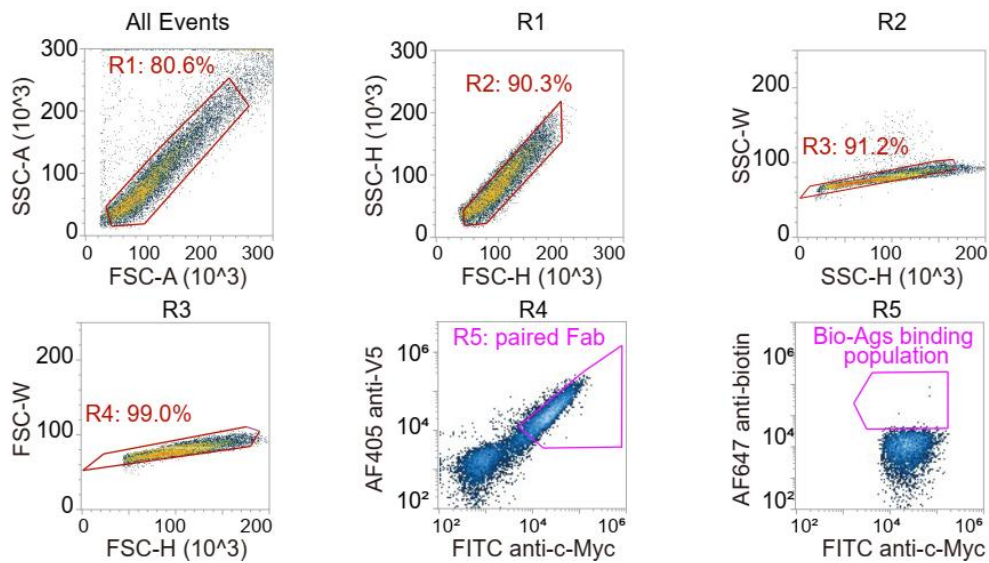

**Figure S2. Flow cytometry gating strategy for yeast display selection.**

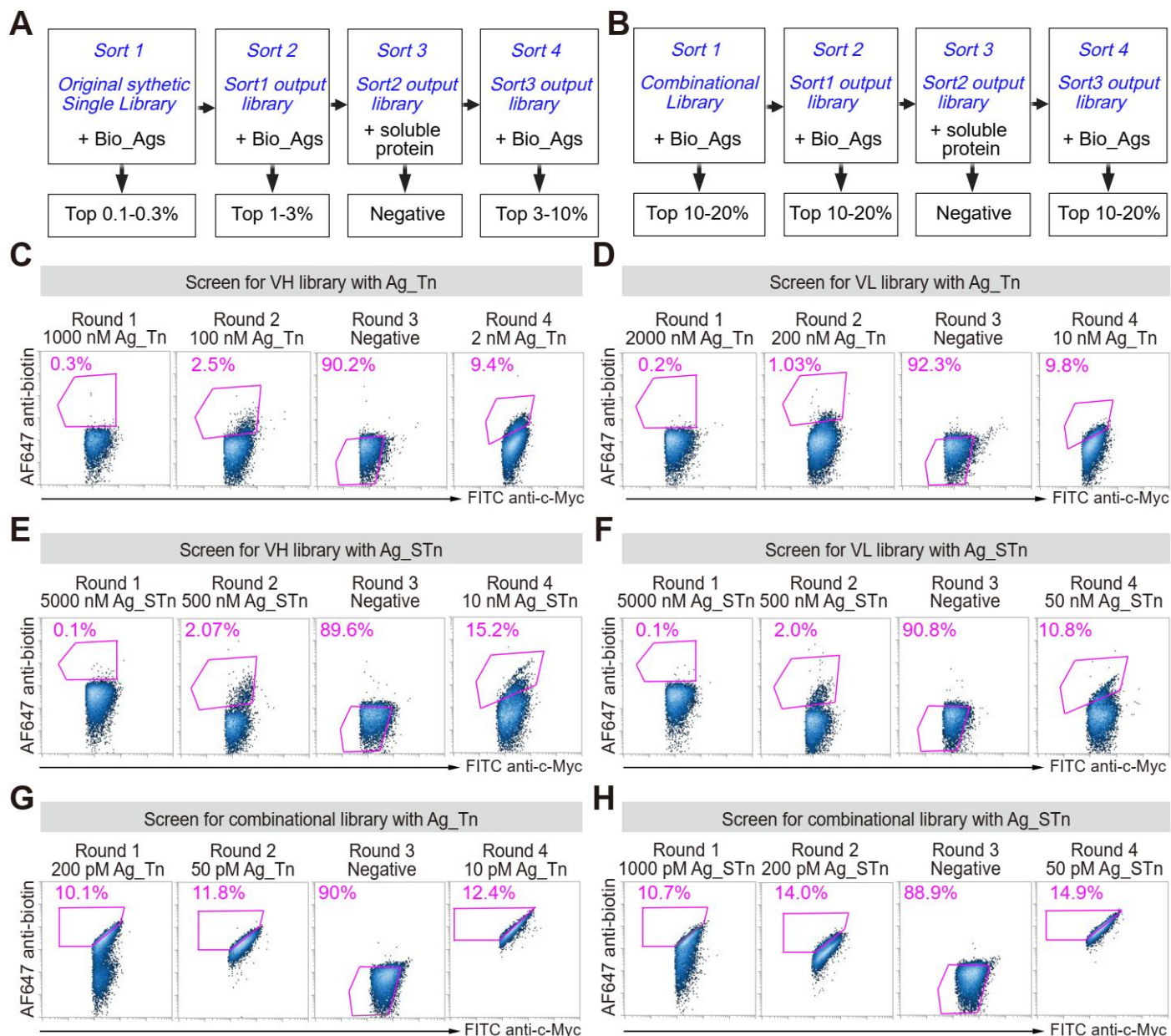

**Figure S3. Yeast display screening and enrichment of antibody variants against MUC1-Tn and MUC1-STn glycoforms.**

**A)** Schematic illustration the library construction and iterative enrichment strategy of single heavy chain (HC) and light chain (LC) variant libraries against Bio-Ags.

**B)** Schematic illustration of the combinatorial HC/LC library and enrichment workflow across four rounds of fluorescence-activated cell sorting (FACS) against Bio-Ags.

**C-D)** Flow cytometry plots from four sequential rounds of FACS screening of the single HC library (**C**) and single LC library (**D**) against MUC1-Tn (Ag\_Tn).

**E-F)** Flow cytometry plots from four sequential rounds of FACS screening of the single HC library (**E**) and single LC library (**F**) against MUC1-STn (Ag\_STn).

**G-H)** Flow cytometry plots from four sequential rounds of FACS screening of the combinatorial HC/LC library against MUC1-Tn (**G**) and MUC1-STn (**H**).

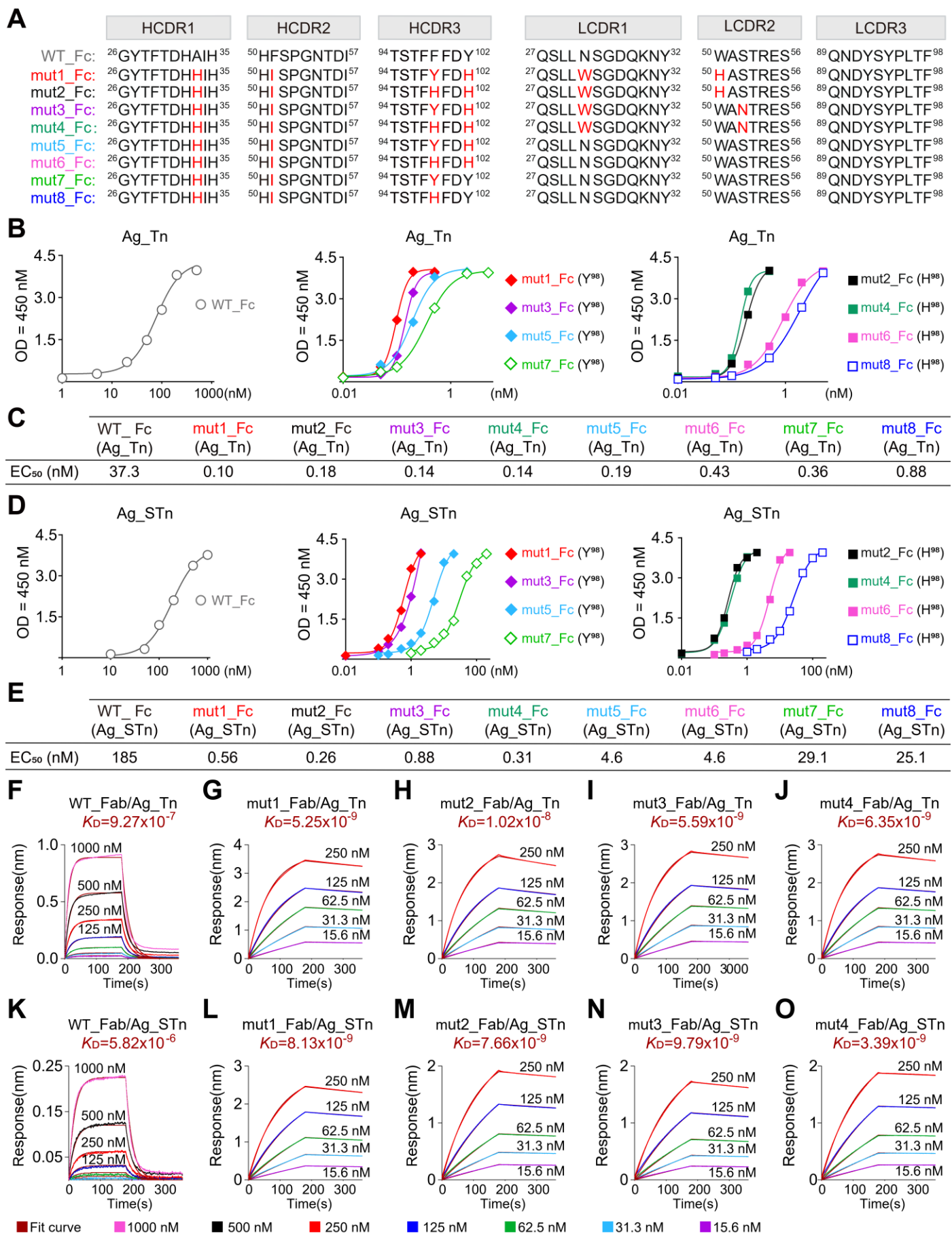

**Figure S4. Enhanced recognition of MUC1-Tn and MUC1-STn glycopeptides by affinity-matured 5E5 variants.**

**A)** CDR sequence alignment of WT and eight affinity-matured variants (mut1–mut8). Mutated residues are highlighted in red bold; wild-type residues at equivalent positions are shown in black bold.

**B)** ELISA binding of WT and affinity-matured scFv-Fc variants to the MUC1-Tn glycopeptide (Ag\_Tn). Variants are grouped according to the residue at HCDR3 position 98 (Y98 or H98).

**C)** EC<sub>50</sub> values derived from the corresponding Ag\_Tn ELISA binding curves shown in (B).

**D)** ELISA binding of WT and affinity-matured scFv-Fc variants to the MUC1-STn glycopeptide (Ag\_STn). Variants are grouped according to the residue at HCDR3 position 98 (Y98 or H98).

**E)** EC<sub>50</sub> values derived from the corresponding Ag\_STn ELISA binding curves shown in (D).

**F-O)** Biolayer interferometry (BLI) analysis of affinity-matured variants. Binding kinetics of parental 5E5 Fab and affinity-matured Mut1–Mut4 Fab fragments against MUC1-Tn (F–J) and MUC1-STn (K–O) glycopeptides, determined using the Octet system. Equilibrium dissociation constants ($K_D$) were calculated from the ratio of dissociation to association rate constants ($k_{off}/k_{on}$).

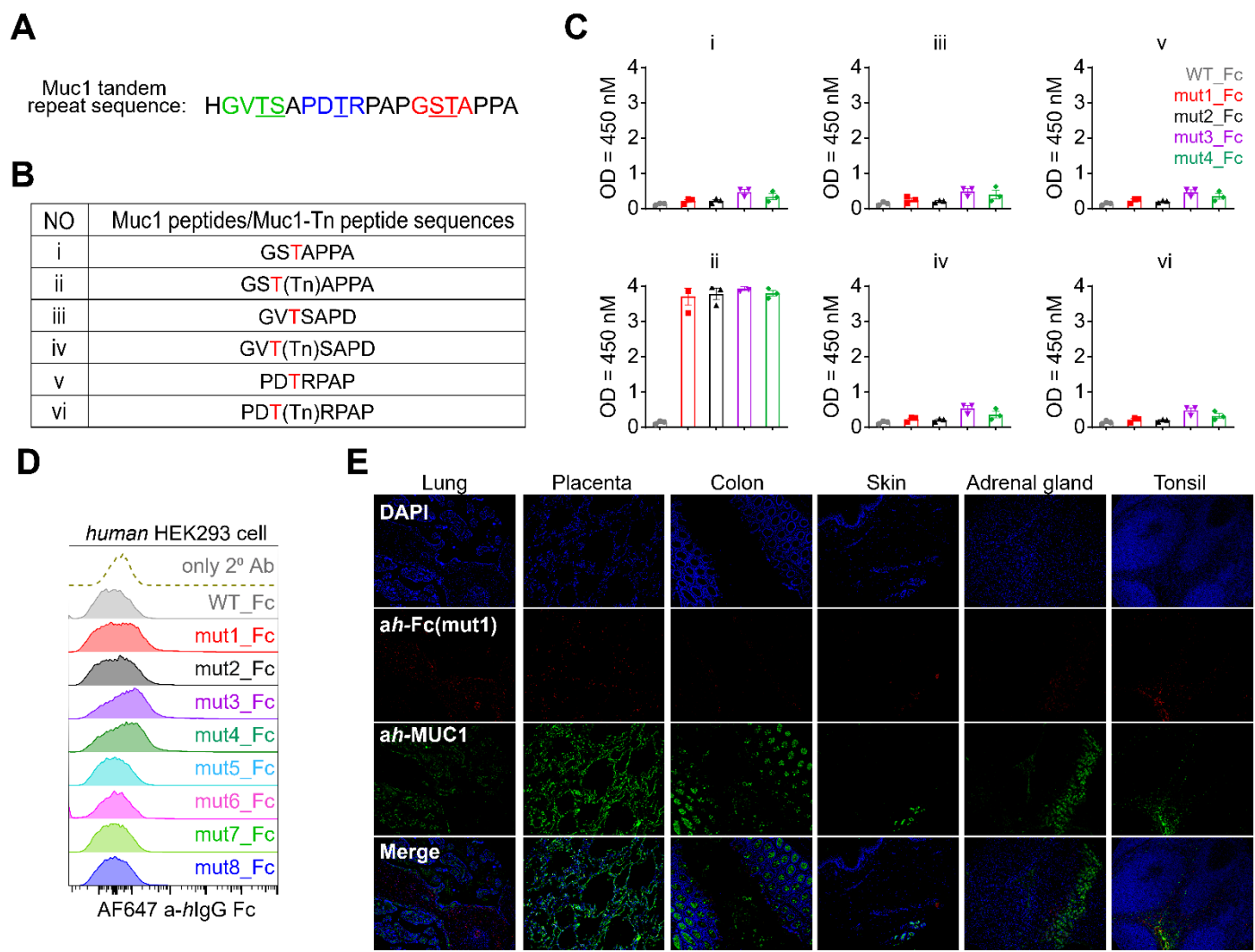

**Figure S5. Epitope specificity profiling and tissue cross-reactivity analysis of affinity-matured 5E5 variants.**

**A)** Schematic representation of the MUC1 variable number of tandem repeats (VNTRs), showing the 20-residue repeat sequence (HGVTSA PDTRPAPGSTAPPA) and three potential epitope regions, GVTS, PDTR, and GSTA.

**B)** Panel of six MUC1-derived antigens used for epitope specificity profiling, comprising three unmodified MUC1 peptides (i, iii, v) spanning the GVTS, PDTR, and GSTA epitope regions and their corresponding Tn-glycosylated counterparts (ii, iv, vi).

**C)** ELISA binding analysis of parental WT 5E5 and affinity-matured variants (mut1–mut4), expressed as Fc-fusion proteins, against the six MUC1 peptide and MUC1-Tn glycopeptide antigens (i–vi) shown in (B). Binding was measured at a fixed antibody concentration of 1 nM.

**D)** Flow cytometric analysis of WT 5E5 and affinity-matured 5E5-Fc fusion variants (mut1–8) binding to normal human HEK293T cells.

**E)** Immunofluorescence analysis of mut1-Fc binding to normal human tissue sections, including lung, placenta, colon, skin, adrenal gland, and tonsil. Bound mut1-Fc was detected with an anti-human Fc antibody (red), endogenous MUC1 was detected with an anti-human MUC1 antibody (green), and nuclei were counterstained with DAPI (blue). Scale bars, 100 μm.

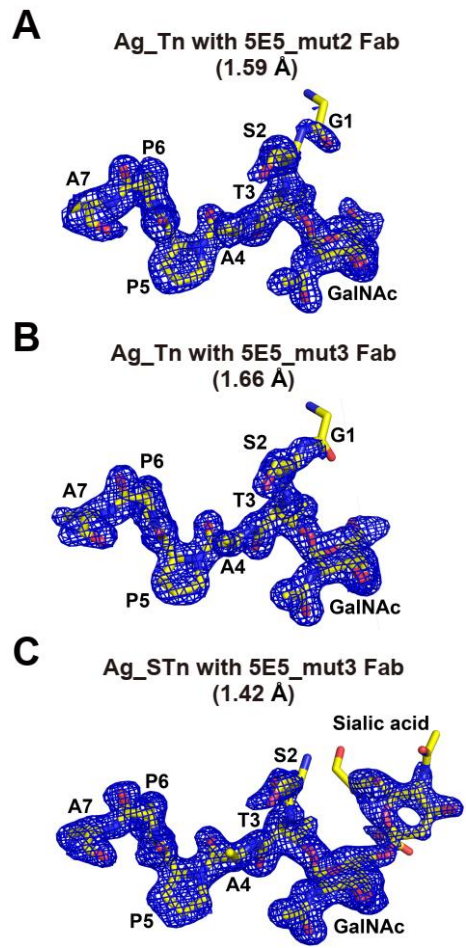

**Figure S6. Electron density maps for peptides Ag\_Tn or Ag\_STn in their complexes with 5E5\_mut2 or 5E5\_mut3 Fab.**

**A)** Crystal structure of 5E5-mut2 Fab in complex with MUC1-Tn glycopeptide (Ag\_Tn), resolved at 1.59 Å. Unbiased 2Fo-Fc electron density omit map (blue mesh, contoured at $1\sigma$) overlaid with the refined glycopeptide structure displayed as sticks. Key peptide residues (A7, P6, P5, A4, T3, S2, G1) and the GalNAc moiety are labelled.

**B)** Crystal structure of 5E5-mut3 Fab in complex with MUC1-Tn glycopeptide (Ag\_Tn), resolved at 1.66 Å. Unbiased 2Fo-Fc electron density omit map (blue mesh, contoured at $1\sigma$) overlaid with the refined glycopeptide structure displayed as sticks. Key peptide residues and the GalNAc moiety are labelled as in **(A)**.

**C)** Crystal structure of 5E5-mut3 Fab in complex with MUC1-STn glycopeptide (Ag\_STn), resolved at 1.42 Å. Unbiased 2Fo-Fc electron density omit map (blue mesh, contoured at $1\sigma$) overlaid with the refined glycopeptide structure displayed as sticks, including the sialic acid moiety. Key peptide residues, GalNAc, and sialic acid are labelled as in **(A)**.

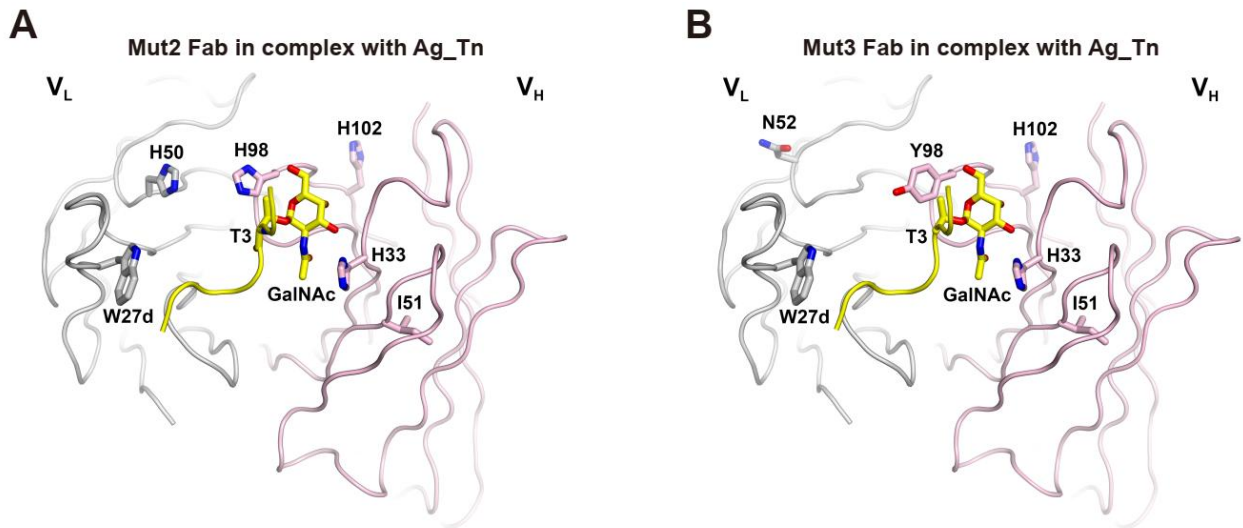

**Figure S7. Mutated residues of the affinity-matured 5E5-mut2 and 5E5-mut3 Fabs as observed in their complex structures with Ag\_Tn**

**A)** Crystal structure of mut2 Fab (V<sub>L</sub>, light grey; V<sub>H</sub>, light pink) in complex with MUC1-Tn glycopeptide.

**B)** Crystal structure of mut3 Fab (V<sub>L</sub>, light grey; V<sub>H</sub>, light pink) in complex with MUC1-Tn glycopeptide, highlighting key residues involved in glycopeptide recognition. HCDR residues H33, I51, H98/Y98, H102, and LCDR residues W27d, H50/N52 are highlighted. The GalNAc moiety and T3 residue are shown as yellow sticks; the differential mutations at position 98 in HCDR3 (Y98 in mut3 *versus* H98 in mut2) and positions in LCDR2 (N52 *versus* His50 in mut2).

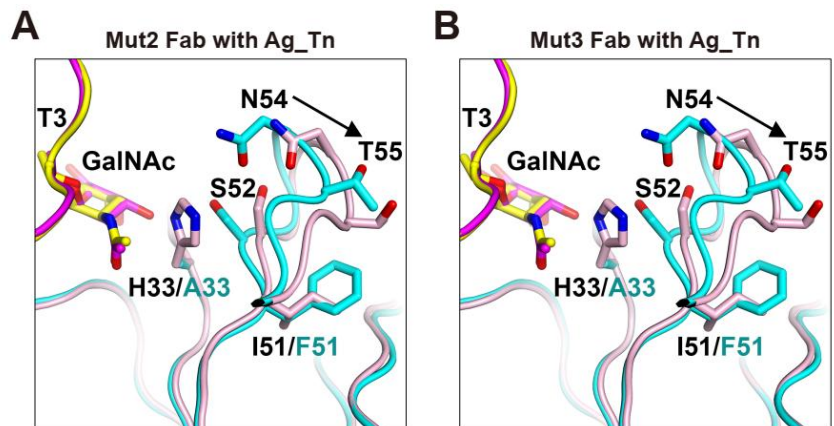

**Figure S8. Structural comparison of 5E5-mut2 or 5E5-mut3 Fabs with 5E5 WT scFv in their complex with Ag\_Tn around mutated HCDR residues 33 and 51.**

**A)** Superimposition of 5E5-mut2 Fab in complex with MUC1-Tn (Ag\_Tn) with WT 5E5 scFv in complex with Ag\_Tn, highlighting conformational changes at HCDR residues H33 and I51.

**B)** Superimposition of 5E5-mut3 Fab in complex with MUC1-Tn (Ag\_Tn) with WT 5E5 scFv in complex with Ag\_Tn, displayed as in (A). For mut2/mut3 complex, H-chain (light pink) was shown with Ag\_Tn peptide in yellow; for WT complex, H-chain (cyan) was shown with Ag\_Tn peptide in magenta. Residue sidechains and the GalNAc moiety are displayed as sticks.

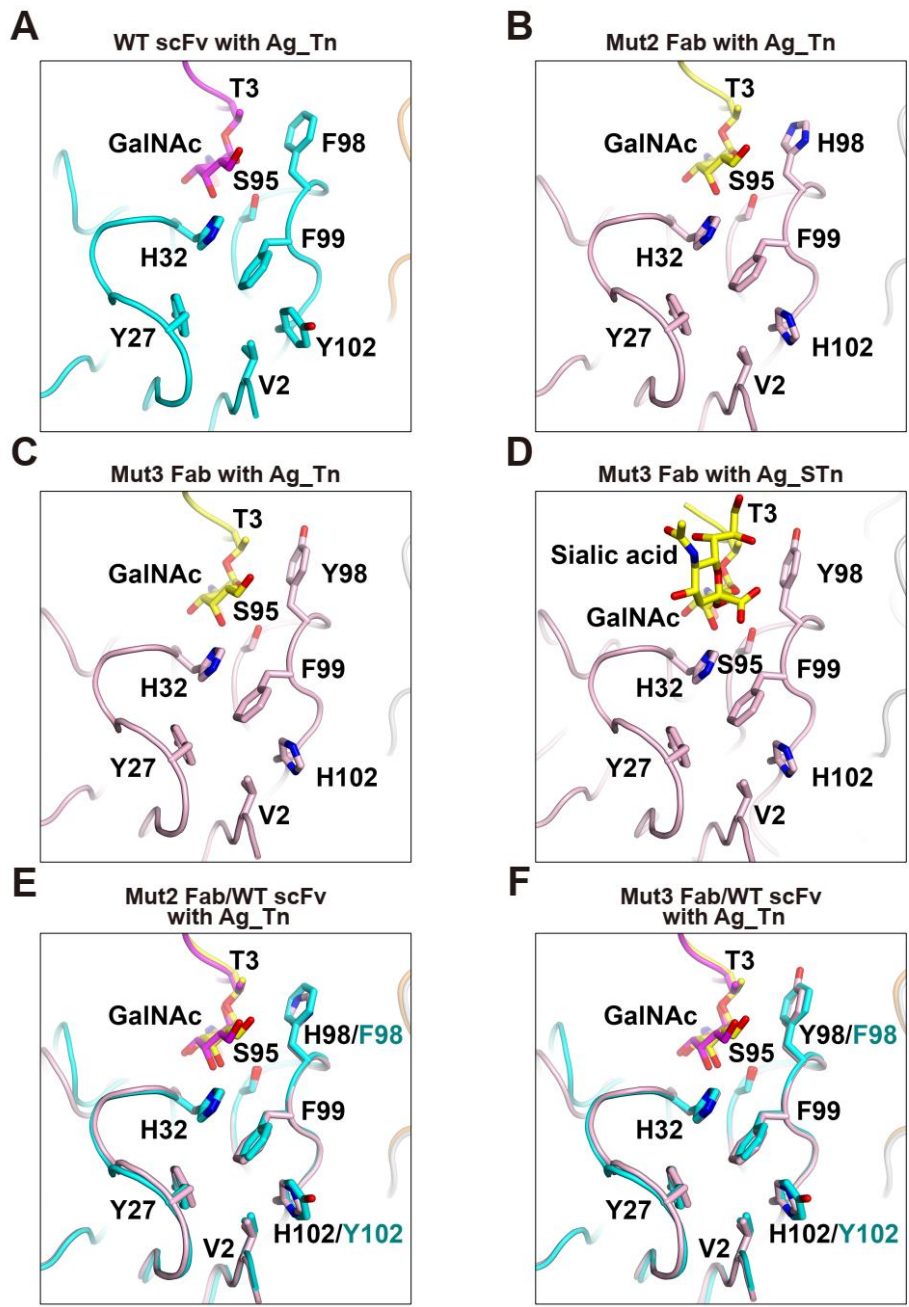

**Figure S9. Structural comparison of 5E5-mut2 or 5E5-mut3 Fabs with 5E5 WT scFv around mutated H-chain residue 102 as observed in their complex with Ag\_Tn or Ag\_STn**

**A)** Close-up view of the HCDR3 binding pocket in WT 5E5 scFv in complex with MUC1-Tn (Ag\_Tn). H-chain (cyan) and L-chain (orange) are shown.

**B)** Close-up view of the HCDR3 binding pocket in 5E5-mut2 Fab in complex with MUC1-Tn (Ag\_Tn). H-chain (light pink) and L-chain (grey) showing the F<sup>98</sup>H and Y<sup>102</sup>H substitutions, which alters the paratope architecture.

**C)** Close-up view of the HCDR3 binding pocket in 5E5-mut3 Fab in complex with MUC1-Tn (Ag\_Tn), displayed as in **B**), showing the Y98 and His102 substitutions.

**D)** Close-up view of the HCDR3 binding pocket in 5E5-mut3 Fab in complex with MUC1-STn (Ag\_STn). H-chain (light pink) and L-chain (grey). The sialic acid and GalNAc moieties are displayed as sticks (yellow).

**E)** Superimposition of 5E5-mut2 Fab (light pink/grey) with WT 5E5 scFv (cyan/orange) in their complexes with Ag\_Tn.

**F)** Superimposition of 5E5-mut3 Fab (light pink/grey) with WT 5E5 scFv (cyan/orange) in their complexes with Ag\_Tn. Key residues Y27, H32, S95, H98, F99 and H102 are highlighted around the GalNAc moiety and Thr3 residue (yellow, displayed as sticks).

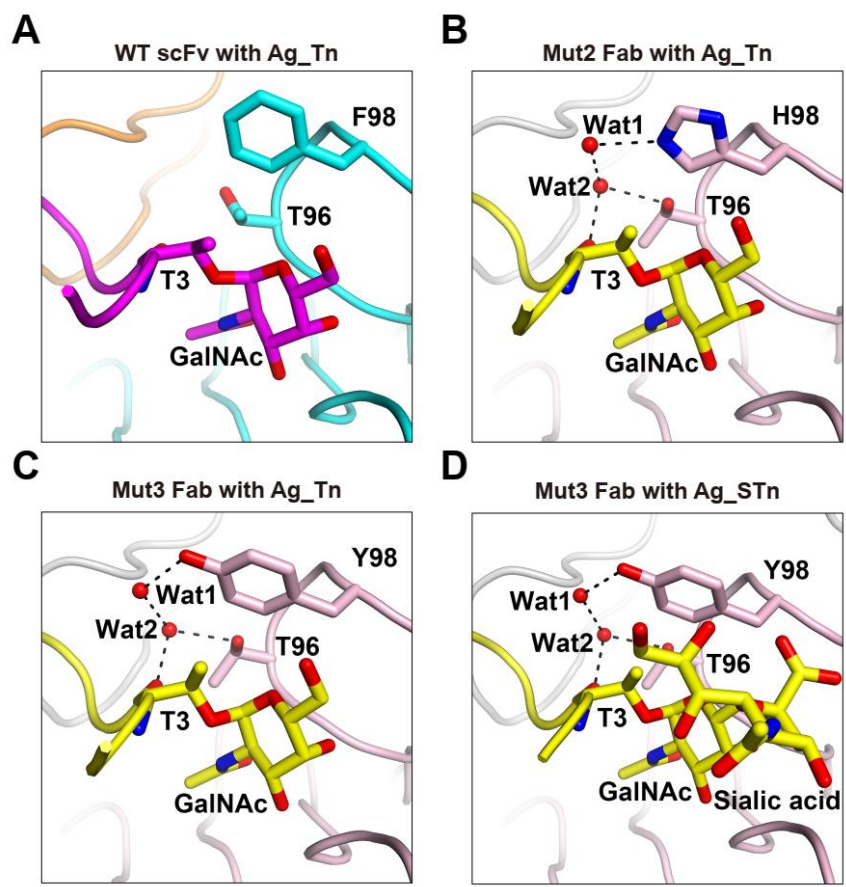

**Figure S10. Structural basis of 5E5\_mut2 or 5E5\_mut3 Fabs with 5E5 WT scFv around mutated H-chain residue 98 as observed in their complex with Ag\_Tn or Ag\_STn**

**A)** Close-up view of the HCDR3 region around F98 of WT 5E5 scFv in complex with MUC1-Tn (Ag\_Tn). The aromatic ring of F98 is positioned adjacent to the GalNAc moiety and Thr3 residue. Residue sidechains and the GalNAc moiety are displayed as sticks.

**B)** Close-up view of the HCDR3 region around H98 of 5E5-mut2 Fab in complex with MUC1-Tn (Ag\_Tn). The F<sup>98</sup>H substitution introduces a polar imidazole ring that enables water-mediated hydrogen bonding interactions (Wat1 and Wat2, shown as black dashed lines) with the GalNAc moiety and T3 residue.

**C)** Close-up view of the HCDR3 region around Y98 of 5E5-mut3 Fab in complex with MUC1-Tn (Ag\_Tn). Water-mediated hydrogen bonding interactions (shown as black dashed lines) connect the aromatic ring and hydroxyl group of Y98 with the GalNAc moiety and T3 residue.

**D)** Close-up view of the HCDR3 region around Y98 of 5E5-mut3 Fab in complex with MUC1-STn (Ag\_STn). Water-mediated hydrogen bonding interactions (shown as black dashed lines) connect the aromatic ring and hydroxyl group of Y98 with the GalNAc moiety, sialic acid, and T3 residue.

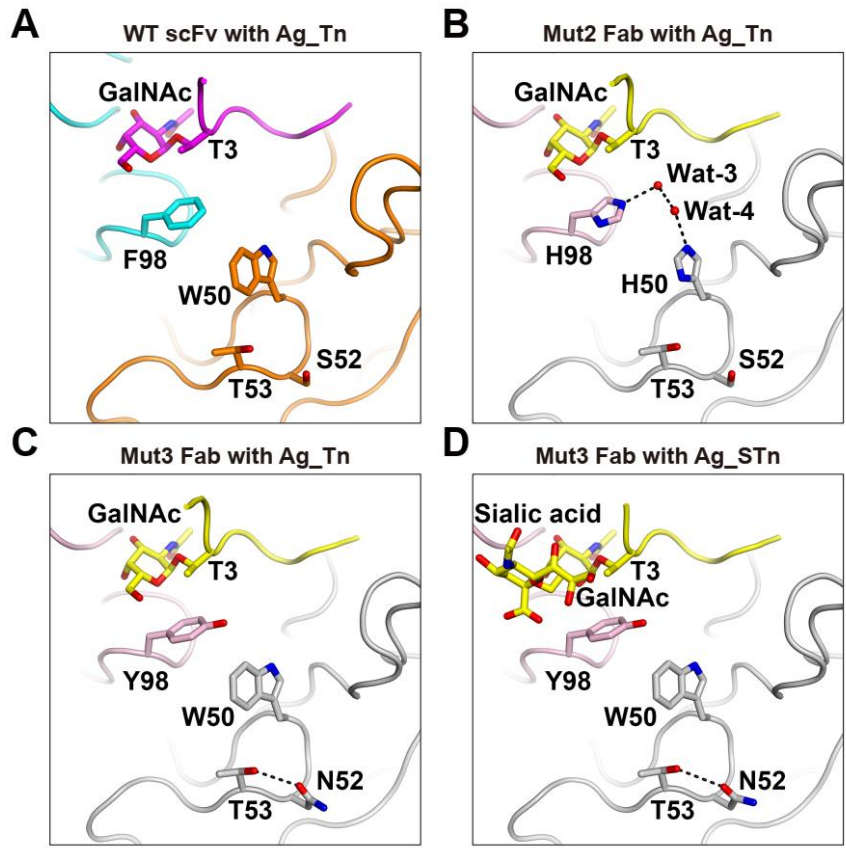

**Figure S11. Structural basis of 5E5\_mut2 and 5E5\_mut3 Fabs with 5E5 WT scFv around L-chain residues 50 and 52 as observed in their complex with Ag\_Tn or Ag\_STn**

**A)** Close-up view of the LCDR2 region around W50 and S52 of WT 5E5 scFv in complex with MUC1-Tn (Ag\_Tn). Residue sidechains and the GalNAc moiety are displayed as sticks.

**B)** Close-up view of the LCDR2 region around H50 and S52 of 5E5-mut2 Fab in complex with MUC1-Tn (Ag\_Tn). The W<sup>50</sup>H substitution introduces a polar imidazole ring that participates in water-mediated hydrogen bonding interactions (Wat-3 and Wat-4, shown as black dashed lines) connecting to the H98 substitution.

**C)** Close-up view of the LCDR2 region around W50 and N52 of 5E5-mut3 Fab in complex with MUC1-Tn (Ag\_Tn). The S52N substitution introduces an intramolecular hydrogen bonding interaction (shown as black dashed lines) between N52 and the sidechain of T53.

**D)** Close-up view of the LCDR2 region around W50 and N52 of 5E5-mut3 Fab in complex with MUC1-STn (Ag\_STn). The S52N substitution maintains the intramolecular hydrogen bonding interaction with T53 (shown as black dashed lines).

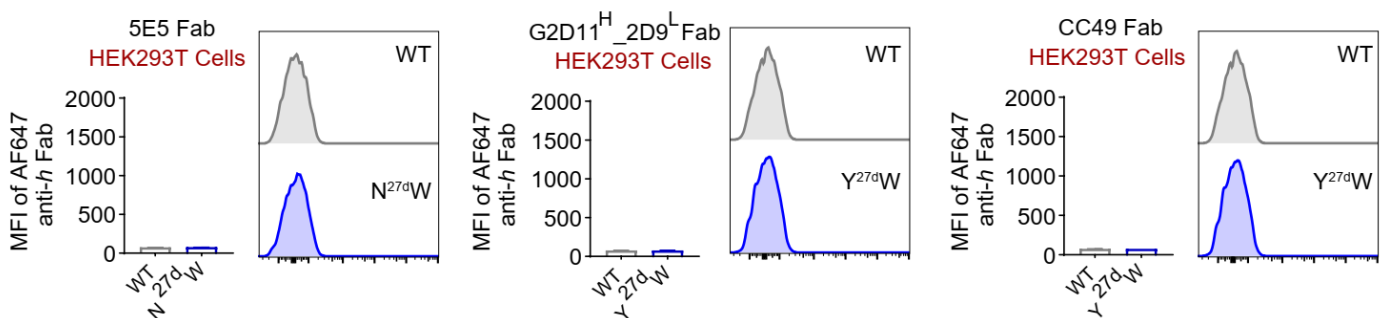

**Figure S12. Affinity-enhancing N/Y<sup>27dW</sup> mutations do not introduce non-specific binding to normal human cells across multiple anti-MUC1 antibody scaffolds**

Flow cytometric analysis of binding by wild-type (WT) and affinity-matured N27d→W (N27dW) Fab fragments from three anti-MUC1 antibodies including 5E5, Chimeric G2D11<sup>H</sup>-2D9<sup>L</sup>, and CC49 to normal human HEK293T cells.

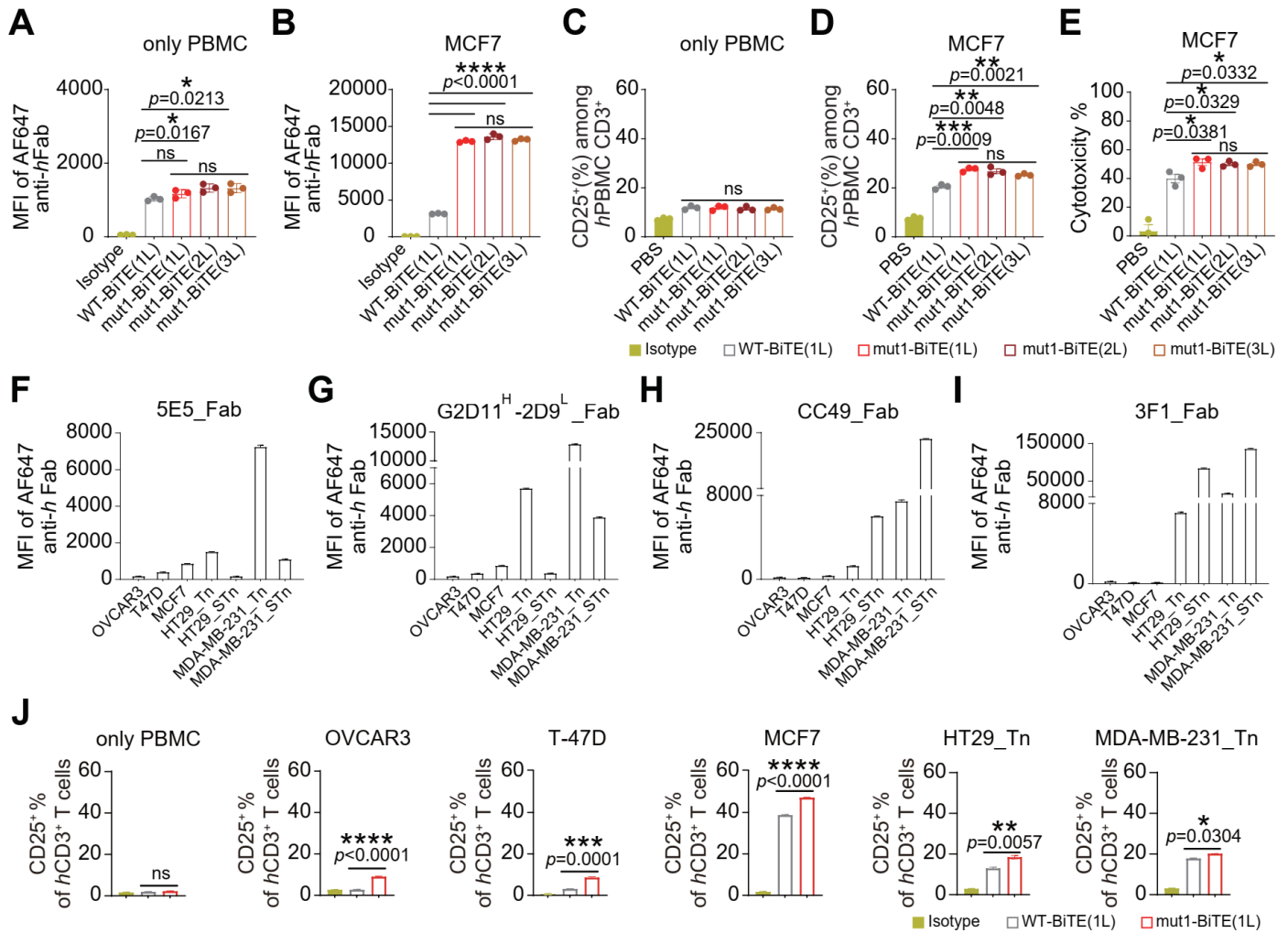

**Figure S13. BiTE linker optimization and T cell activation across MUC1-Tn/STn-expressing cancer cell lines**

**A-B** Flow cytometric analysis of BiTE binding with varying G4S linker lengths (1L, 2L, 3L) to normal human PBMCs (**A**) and MCF7 breast cancer cells (**B**). Mean fluorescence intensity (MFI) of AF647-labelled BiTE fragments are shown with statistical significance indicated.

**C-D** Flow cytometric assessment of T cell activation, quantified as the percentage of CD25<sup>+</sup> cells among CD3<sup>+</sup> T cells, induced by BiTE molecules with different G4S linker lengths in PBMCs alone (**C**) and in co-culture of primary human with MCF7 cells (**D**).

**E** Cytolytic efficacy of BiTE molecules with varying G4S linker lengths measured by live-cell analysis (Incucyte CX3) in co-culture of primary human PBMCs and MCF7 breast cancer cells at a 5:1 effector-to-target ratio.

**F-I** Flow cytometry binding analysis of Fab fragments from four anti-Tn/STn antibodies including 5E5 (**F**), G2D11<sup>H</sup>-2D9<sup>L</sup> (**G**), CC49 (**H**), and 3F1 (**I**) against a panel of cancer cell lines (OVCAR3, T47D, MCF7 HT29\_Tn, HT29\_STn, MDA-MB-231\_Tn, and MDA-MB-231 cells). Binding was detected using an AF647-conjugated anti-human Fab secondary antibody and quantified as mean fluorescence intensity (MFI).

**J** T cell activation induced by PBS, WT-BiTE, and mut1-BiTE in the presence of the indicated cancer cell lines (OVCAR3, T-47D, MCF7, HT29\_Tn, MB-MDA-231\_Tn) or PBMCs alone, quantified as the percentage of CD25<sup>+</sup> cells among CD3<sup>+</sup> T cells by flow cytometry. Statistical comparisons between WT-BiTE and mut1-BiTE are indicated. Data represent mean ± SEM of $n = 3$ independent experiments. ns, not significant; \*, $p < 0.05$; \*\*, $p < 0.01$; \*\*\*\*, $p < 0.0001$.

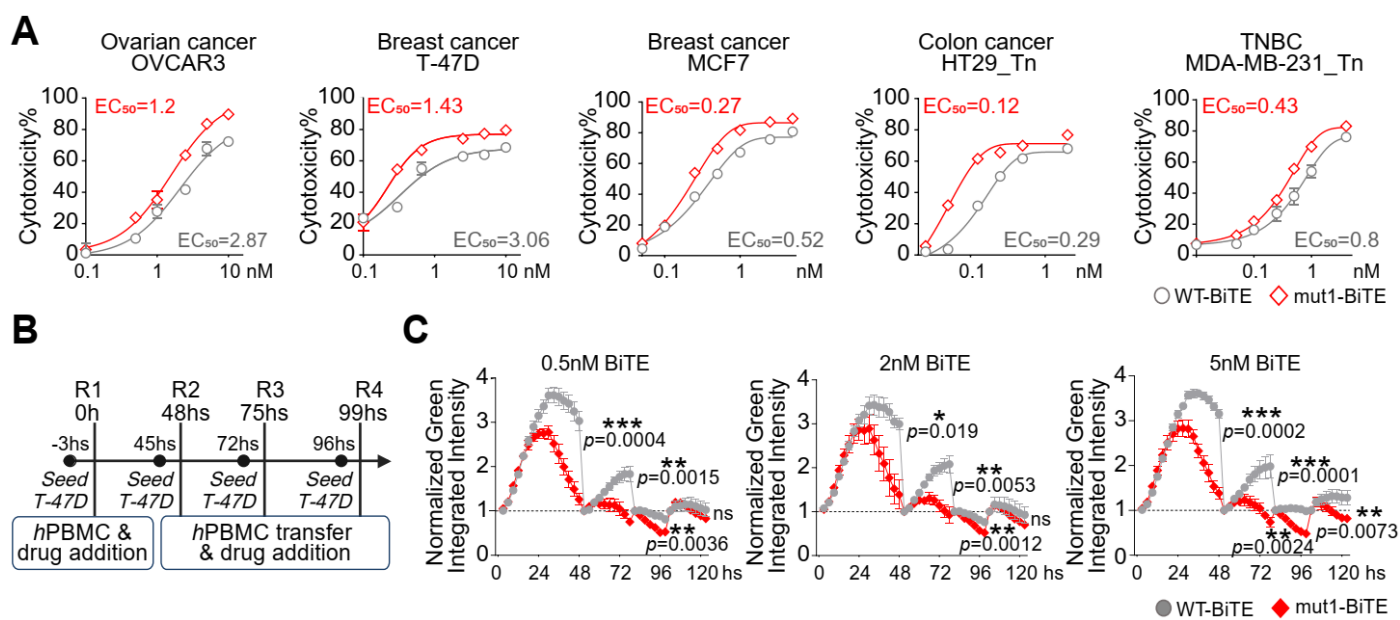

**Figure S14. Bispecific T cell engagers derived from affinity-matured 5E5 variants display enhanced cytolytic activity**

**A)** Dose-response curves for cytotoxicity of WT-BiTE (grey circles) and mut1-BiTE (red diamonds) against five MUC1-Tn/STn-expressing cancer cell lines: ovarian cancer OVCAR3, breast cancers T-47D and MCF7, colon cancer HT29\_Tn, and triple-negative breast cancer MDA-MB-231\_Tn. Cytotoxicity (%) is plotted against BiTE concentration (0.1–10 nM). $EC_{50}$ values are indicated for each cell line. Data represent mean ± SEM of $n = 3$ independent experiments.

**B)** Schematic representation of the serial killing assay design. T-47D tumor cells were seeded and co-cultured with primary human PBMCs and BiTE constructs. Four sequential rounds (R1–R4) of tumor cell killing were performed, with PBMC transfer and fresh BiTE addition at 48-hour, 72-hour, 96-hour to assess repeated killing capacity.

**C)** Serial killing effect of WT-BiTE (grey) and mut1-BiTE (red) against T-47D cells at three BiTE concentrations (0.5 nM, 2 nM, 5 nM). Normalized green integrated intensity (reflecting viable tumor cell) is plotted over time (0–120 hours). Statistical comparisons between WT-BiTE and mut1-BiTE are indicated: ns, not significant; \*, $p < 0.05$; \*\*, $p < 0.01$; \*\*\*\*, $p < 0.0001$. Data represent mean ± SEM of $n = 3$ independent experiments.

WT-BiTE and e4-BiTE triggered cytotoxicity via co-culture of primary *human* normal PBMC (donor#2) and indicated cancer cell lines for 2–3 days and with E:T ratios at 5:1.

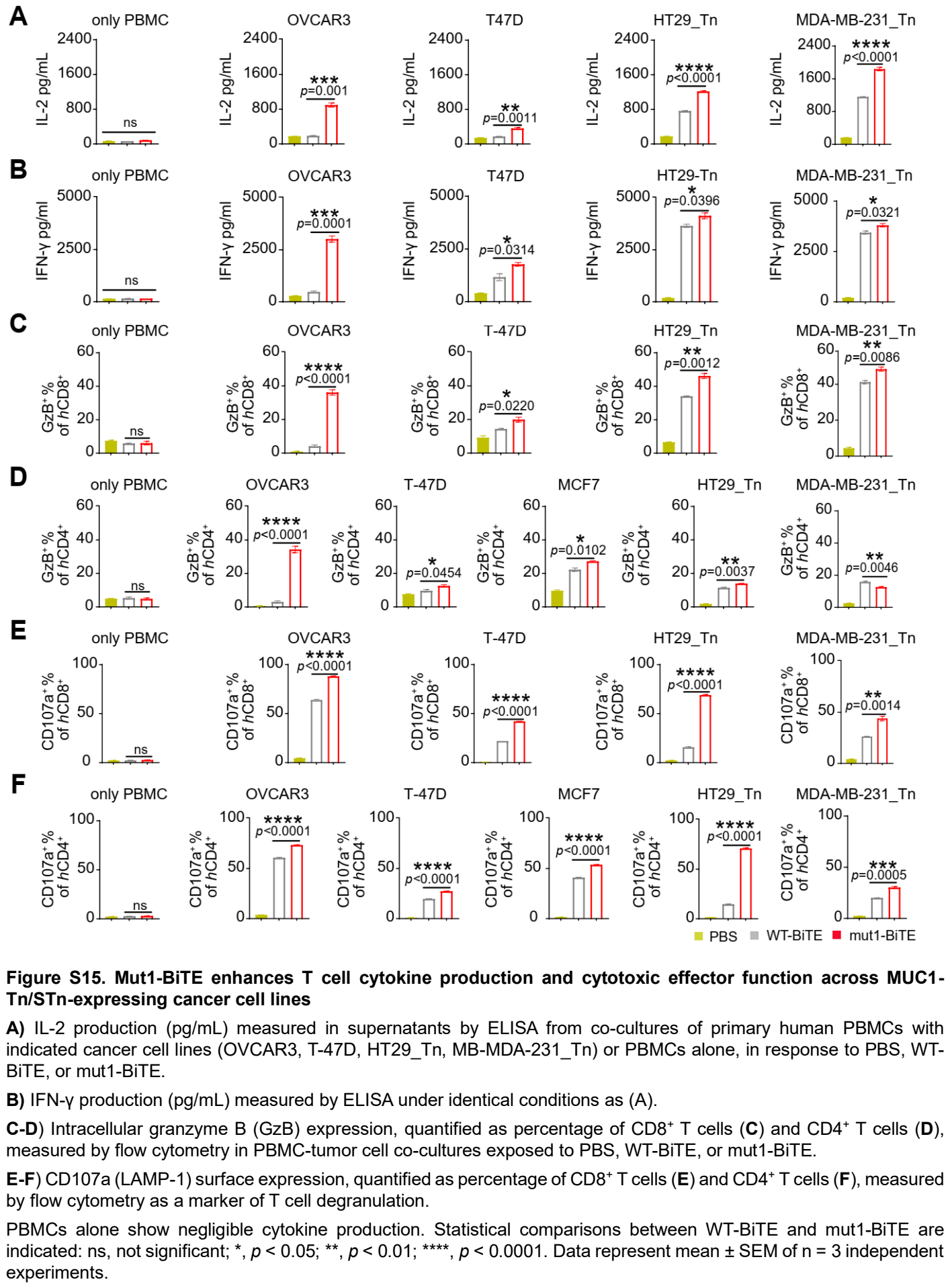

Figure S15. Mut1-BiTE enhances T cell cytokine production and cytotoxic effector function across MUC1- Tn/STn-expressing cancer cell lines

A) IL-2 production (pg/mL) measured in supernatants by ELISA from co-cultures of primary human PBMCs with indicated cancer cell lines (OVCAR3, T-47D, HT29\_Tn, MB-MDA-231\_Tn) or PBMCs alone, in response to PBS, WT-BiTE, or mut1-BiTE.

B) IFN-γ production (pg/mL) measured by ELISA under identical conditions as (A).

C-D) Intracellular granzyme B (GzB) expression, quantified as percentage of CD8+ T cells (C) and CD4 + T cells (D), measured by flow cytometry in PBMC-tumor cell co-cultures exposed to PBS, WT-BiTE, or mut1-BiTE.

E-F) CD107a (LAMP-1) surface expression, quantified as percentage of CD8+ T cells (E) and CD4 + T cells (F), measured by flow cytometry as a marker of T cell degranulation.

PBMCs alone show negligible cytokine production. Statistical comparisons between WT-BiTE and mut1-BiTE are indicated: ns, not significant; \*, p < 0.05; \*\*, p < 0.01; \*\*\*\*, p < 0.0001. Data represent mean ± SEM of n = 3 independent experiments.

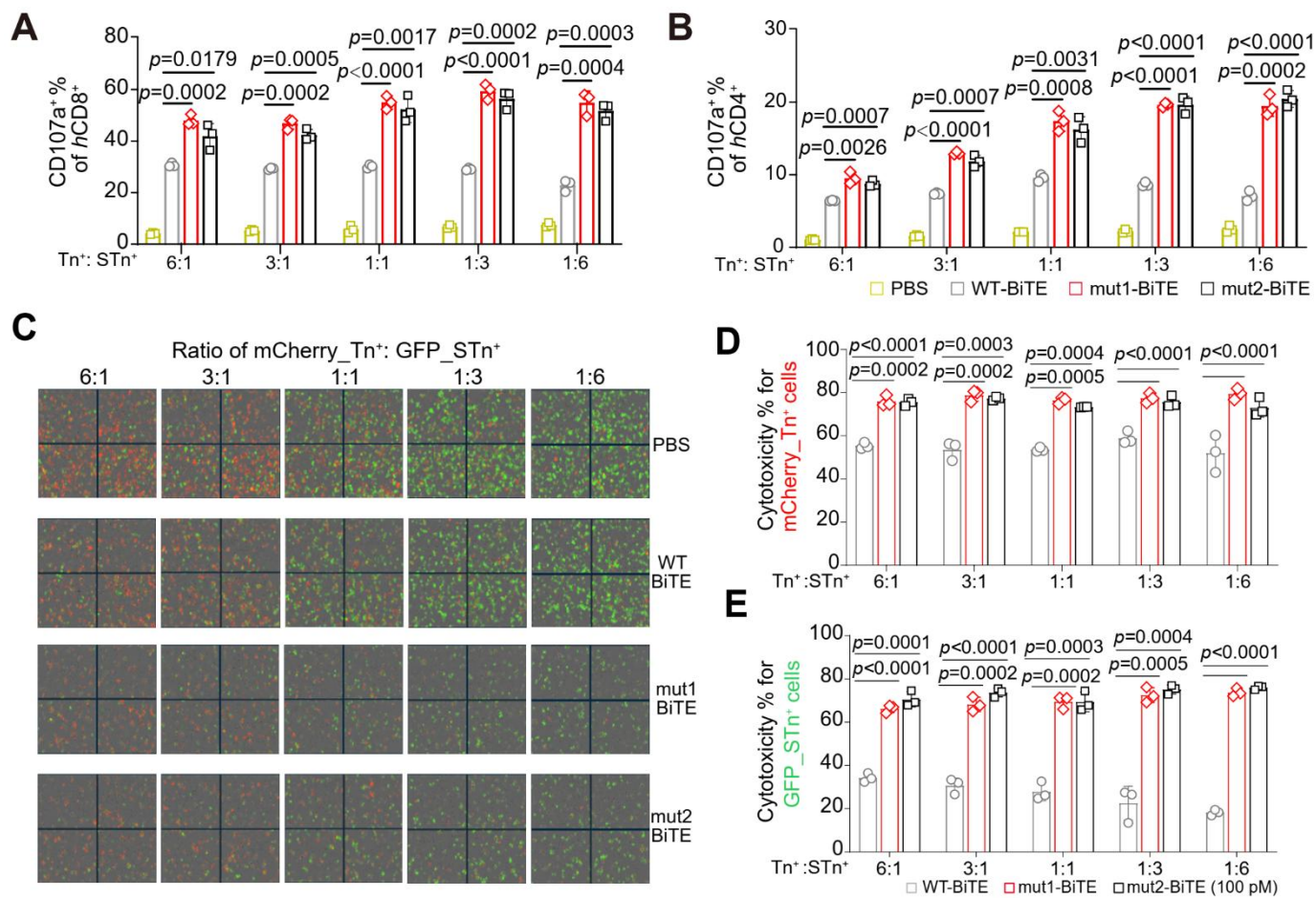

**Figure S16. Characterization of BiTE activity against heterogeneous Tn/STn-expressing tumor cell populations.**

**A, B** CD107a expression in CD8<sup>+</sup> (**A**) and CD4<sup>+</sup> (**B**) T cells following co-culture with mixed HT29\_Tn<sup>+</sup> and HT29\_STn<sup>+</sup> tumor cells at the indicated Tn<sup>+</sup>: STn<sup>+</sup> ratios in the presence of PBS, WT-BiTE, mut1-BiTE, or mut2-BiTE.

**C** Representative fluorescence images of mixed HT29\_Tn<sup>+</sup> and HT29\_STn<sup>+</sup> cells at the indicated Tn<sup>+</sup>: STn<sup>+</sup> ratios following treatment with PBS, WT-BiTE, mut1-BiTE, or mut2-BiTE (100 pM). HT29\_Tn<sup>+</sup> and HT29\_STn<sup>+</sup> cells were distinguished by mCherry and GFP expression, respectively.

**D, E** Quantification of cytotoxicity against mCherry<sup>+</sup> Tn<sup>+</sup> (**D**) and GFP<sup>+</sup> STn<sup>+</sup> (**E**) tumor cells at the indicated Tn<sup>+</sup>: STn<sup>+</sup> ratios following treatment with WT-BiTE, mut1-BiTE, or mut2-BiTE (100 pM).

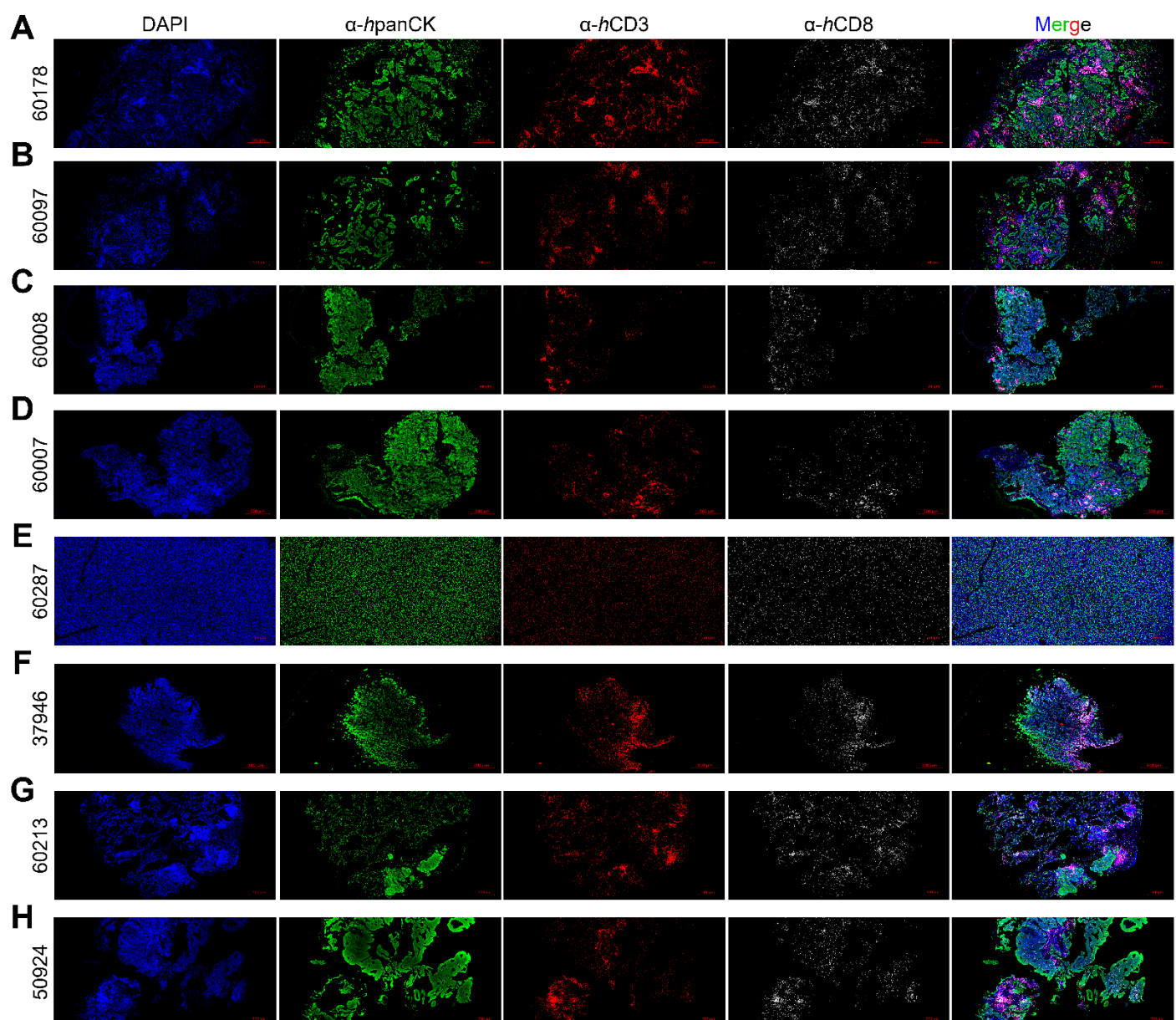

**Figure S17. Immunofluorescence characterization of T-cell infiltration in BiTE-responsive patient-derived tumor samples.**

**A–H)** Representative immunofluorescence images of BiTE-responsive patient-derived tumor samples 60178, 60097, 60007, 60008, 37946, 60287, 50924, and 60213, respectively. Sections were stained with DAPI (blue), pan-cytokeratin (panCK; green), CD3 (red), and CD8 (white), with merged images shown in the rightmost column. Scale bars, 500 μm.

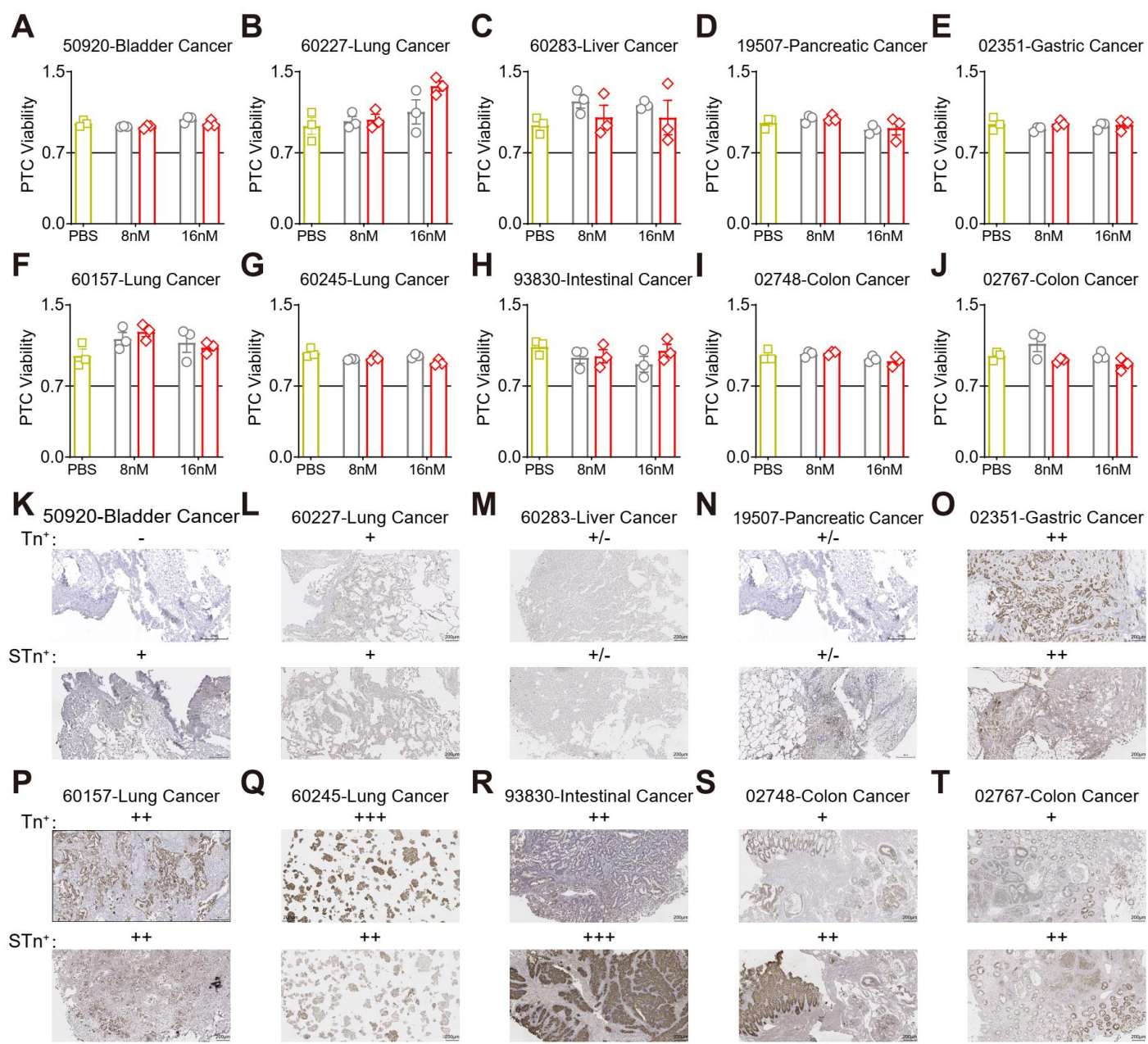

**Figure S18. Representative non-responsive PTC samples to WT and affinity-matured mut1 BiTEs**

**A–J** Viability of PTCs derived from ten independent patient tumor samples including bladder (50920), lung (60227, 60157, 60245), liver (60283), pancreatic (19507), gastric (02351), intestinal (93830), and colon (02748, 02767) cancers, following treatment with PBS, or 5E5 WT-BiTE or mut1-BiTE at 8 nM and 16 nM. Data are presented as mean ± SEM.

**K–T** Representative immunohistochemical staining for Tn and STn antigen expression in the corresponding PTC samples shown in (A–J), scored semi-quantitatively as negative (–), weak/equivocal (+/–), weak (+), moderate (++), or strong (+++) staining. Scale bars, 200 μm.

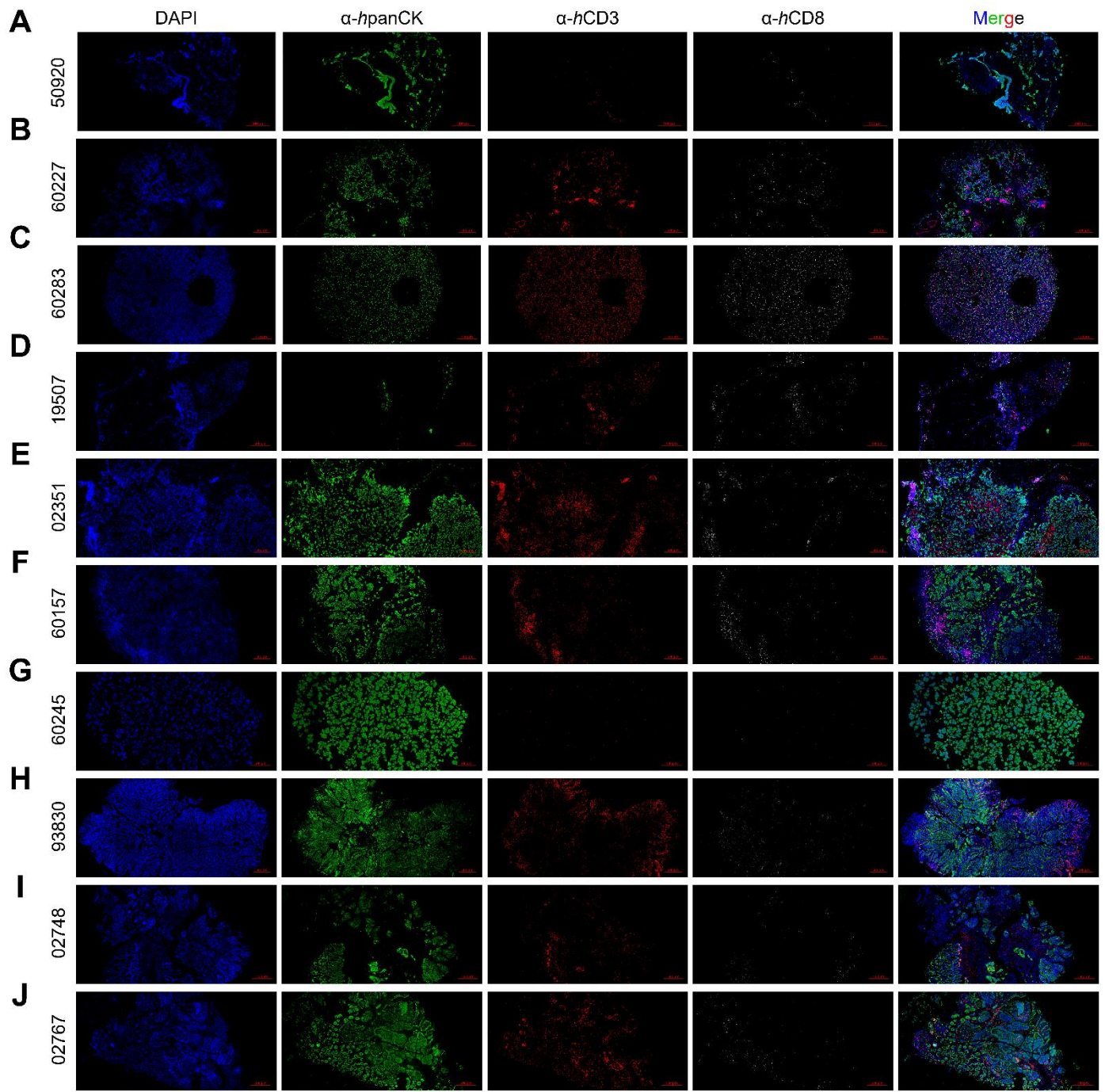

**Figure S19. Immunofluorescence characterization of immune-cell infiltration for nonresponsive PTCs**

**A–J)** Representative immunofluorescence images of patient-derived tumor tissues stained with DAPI (blue), pan-cytokeratin (panCK; green), CD3 (red), and CD8 (white), with merged images shown in the rightmost column. Sample IDs are indicated on the left. Scale bars, 500 μm.

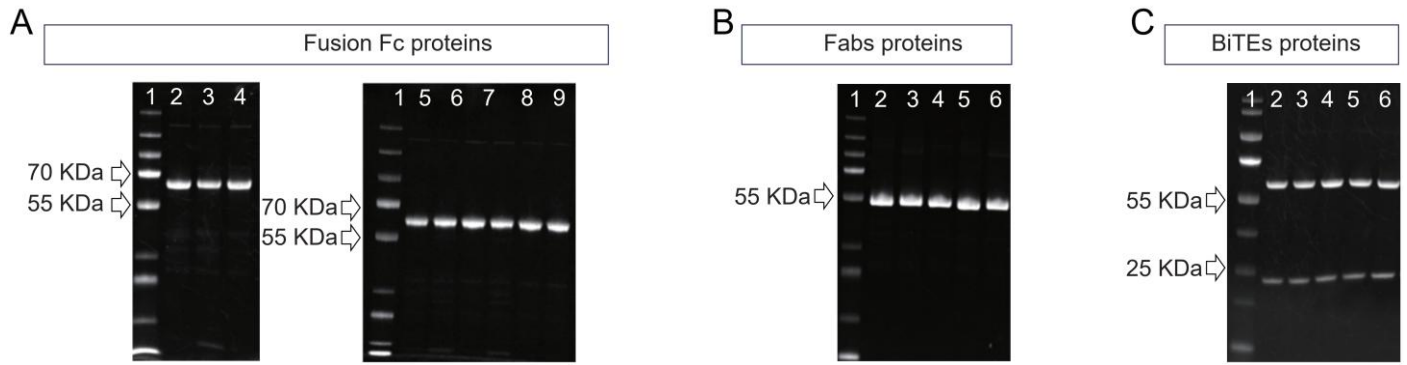

**Figure S20. SDS-PAGE analysis confirming purity of affinity-matured 5E5 Fc-fusion, Fab, and BiTE proteins**

**(A)** Reducing SDS-PAGE of eleven affinity-matured 5E5-Fc fusion proteins. Lane 1, molecular weight marker; lanes 2–9, individual 5E5 WT and mut1-8 Fc-fusion proteins; Protein bands appear at expected molecular weights (~60 kDa for intact monomeric Fc-fusion).

**(B)** Non-reducing SDS-PAGE of five 5E5 Fab proteins. Lane 1, molecular weight marker; lanes 2–6, individual 5E5 WT and mut1-4 Fab proteins. Protein bands appear at the expected molecular weight (~55 kDa) for intact Fab fragments.

**(C)** Reducing SDS-PAGE of six 5E5 BiTE proteins. Lane 1, molecular weight marker; lane 2, WT-BiTE(1L); lane 3, mut1-BiTE(1L); lane 4, mut2-BiTE(1L); lane 5, mut1-BiTE(2L); lane 6, mut1-BiTE(3L). Protein bands appear at expected molecular weights (~55 kDa for HC anti-CD3 scFv and ~25 kDa for LC).

**Table S1. X-ray data collection and refinement statistics**

| Data set | 5E5 scFv<br>Ag_Tn | 5E5-mut2 Fab<br>Ag_Tn | 5E5-mut3 Fab<br>Ag_Tn | 5E5-mut3 Fab<br>Ag_STn |
| --- | --- | --- | --- | --- |
| **Data Collection** |  |  |  |  |
| Wavelength (Å) | 0.97946 | 0.97946 | 0.97946 | 0.97946 |
| Space group | P2<sub>1</sub> | C2 | C2 | C2 |
| Unit cell (Å) | *a* = 67.3<br>*b* = 147.5<br>*c* = 82.2 | *a* = 94.4<br>*b* = 60.7<br>*c* = 80.3 | *a* = 94.7<br>*b* = 60.8<br>*c* = 79.9 | *a* = 94.7<br>*b* = 60.7<br>*c* = 80.3 |
| Angle (°) | 90, 101.4, 90 | 90, 93.2, 90 | 90, 93.2, 90 | 90, 92.9, 90 |
| Resolution (Å)<sup>a</sup> | 38.90-2.35<br>(2.46-2.35) | 40.07-1.60<br>(1.63-1.60) | 39.88-1.66<br>(1.69-1.66) | 31.98-1.42<br>(1.44-1.42) |
| Unique reflections<sup>a</sup> | 60,187 (4,008) | 56,190 (2,147) | 52,734 (2,387) | 83,632 (3,283) |
| Redundancy<sup>a</sup> | 3.2 (2.9) | 3.2 (2.1) | 6.0 (3.9) | 5.8 (2.7) |
| Average I/σ(I)<sup>a</sup> | 8.2 (0.8) | 21.6 (1.5) | 15.2 (1.1) | 32.4 (1.5) |
| Completeness (%)<sup>a</sup> | 95.8 (95.5) | 93.1 (72.1) | 98.0 (89.8) | 96.8 (75.5) |
| *R*<sub>sym</sub><sup>a,b</sup> | 0.16 (0.99) | 0.08 (0.60) | 0.14 (>1) | 0.10 (0.71) |
| *R*<sub>pim</sub><sup>a,b</sup> | 0.10 (0.66) | 0.05 (0.50) | 0.06 (0.63) | 0.04 (0.42) |
| CC<sub>1/2</sub><sup>a</sup> | 0.985 (0.473) | 0.996 (0.545) | 0.991 (0.511) | 0.996 (0.713) |
| No. scFv or Fab per ASU<sup>c</sup> | 6 | 1 | 1 | 1 |
| **Refinement** |  |  |  |  |
| Resolution (Å) | 38.90-2.35 | 40.07-1.60 | 39.88-1.66 | 31.98-1.42 |
| Reflections in refinement | 60,027 | 56,168 | 52,656 | 83,517 |
| Refined residues | 1,419 | 454 | 453 | 455 |
| Refined waters | 296 | 526 | 552 | 522 |
| *R*<sub>cryst</sub><sup>d</sup> | 0.214 | 0.150 | 0.145 | 0.164 |
| *R*<sub>free</sub><sup>e</sup> | 0.259 | 0.182 | 0.181 | 0.188 |
| *B*-values (Å<sup>2</sup>) |  |  |  |  |
| scFv or Fab | 54 | 22 | 19 | 24 |
| peptide | 62 | 31 | 28 | 38 |
| Water | 49 | 33 | 30 | 33 |
| Wilson *B*-values (Å<sup>2</sup>) | 43 | 17 | 16 | 16 |
| Ramachandran values (%)<sup>f</sup> | 96.3, 0 | 96.8, 0 | 97.5, 0 | 97.5, 0 |
| r.m.s.d. bond (Å) | 0.004 | 0.008 | 0.010 | 0.007 |
| r.m.s.d. angle (deg.) | 0.79 | 1.03 | 1.11 | 1.00 |
| PDB codes | 38SP | 38SQ | 38SR | 38SS |

<sup>a</sup> Parentheses denote outer-shell statistics.

<sup>b</sup> $R_{\text{sym}} = \sum_{hkl} \sum_i |I_{hkl,i} - \langle I_{hkl} \rangle| / \sum_{hkl} \sum_i I_{hkl,i}$ and $R_{\text{pim}} = \sum_{hkl} [1/(N-1)]^{1/2} \sum_i |I_{hkl,i} - \langle I_{hkl} \rangle| / \sum_{hkl} \sum_i I_{hkl,i}$, where $I_{hkl,i}$ is the scaled intensity of the *i*<sup>th</sup> measurement of reflection *h, k, l*, $\langle I_{hkl} \rangle$ is the average intensity for that reflection, and *N* is the redundancy. $R_{\text{pim}} = \sum_{hkl} (1/(n-1))^{1/2} \sum_i |I_{hkl,i} - \langle I_{hkl} \rangle| / \sum_{hkl} \sum_i I_{hkl,i}$, where *n* is the redundancy

<sup>c</sup> No. scFv or Fab refers to number of scFv or Fab per asymmetric unit (ASU).

<sup>d</sup> $R_{\text{cryst}} = \sum_{hkl} |F_o - F_c| / \sum_{hkl} |F_o|$, where *F<sub>o</sub>* and *F<sub>c</sub>* are the observed and calculated structure factors.

<sup>e</sup> *R*<sub>free</sub> was calculated as for *R*<sub>cryst</sub>, but on 5% of data excluded before refinement.

<sup>f</sup> Percentage of residues in the favored and outliers regions analyzed by MolProbity.

**Table S2: Quantification of tumor-infiltrating T-cell composition across eighteen PTC specimens**

| PTCs ID | DAPI<sup>+</sup><br>Cell Count | CD8<sup>+</sup> DAPI<sup>+</sup><br>Cell Count | CD3<sup>+</sup> DAPI<sup>+</sup><br>Cell Count | CD8<sup>+</sup>/DAPI<sup>+</sup><br>(%) | CD3<sup>+</sup>/DAPI<sup>+</sup><br>(%) | CD8<sup>+</sup>/CD3<sup>+</sup><br>(%) |
| --- | --- | --- | --- | --- | --- | --- |
| 60178 | 10983 | 3084 | 4183 | 28.08 | 38.09 | 73.73 |
| 37946 | 2023 | 1389 | 1807 | 68.66 | 89.32 | 76.87 |
| 60097 | 8263 | 1976 | 3221 | 23.91 | 38.98 | 61.35 |
| 60007 | 5780 | 1283 | 2294 | 22.20 | 39.69 | 55.93 |
| 60008 | 3296 | 1190 | 1414 | 36.10 | 42.90 | 84.16 |
| 60287 | 14160 | 4058 | 11043 | 28.66 | 77.99 | 36.75 |
| 50924 | 4406 | 1947 | 2255 | 44.19 | 51.18 | 86.34 |
| 60213 | 5394 | 2916 | 4115 | 54.06 | 76.29 | 70.86 |
| 50920 | 2831 | 72 | 145 | 2.54 | 5.12 | 49.66 |
| 60227 | 6360 | 584 | 1602 | 9.18 | 25.19 | 36.45 |
| 60283 | 11926 | 2075 | 4684 | 17.40 | 39.28 | 44.30 |
| 19507 | 4165 | 718 | 1334 | 17.24 | 32.03 | 53.82 |
| 02351 | 11655 | 831 | 2885 | 7.13 | 24.75 | 28.80 |
| 60157 | 14160 | 888 | 2271 | 6.27 | 16.04 | 39.10 |
| 60245 | 5973 | 5 | 115 | 0.08 | 1.93 | 4.35 |
| 93830 | 8343 | 309 | 3656 | 3.70 | 43.82 | 8.45 |
| 02748 | 8286 | 112 | 606 | 1.35 | 7.31 | 18.48 |
| 02767 | 9360 | 92 | 1599 | 0.98 | 17.08 | 5.75 |

CD3<sup>+</sup> and CD8<sup>+</sup> cells were counted only when the corresponding immunofluorescence signal was associated with a DAPI<sup>+</sup> nucleus. Percentages of CD3<sup>+</sup> and CD8<sup>+</sup> cells were calculated relative to the total number of DAPI<sup>+</sup> cells. The CD8<sup>+</sup>/CD3<sup>+</sup> ratio represents the proportion of CD8<sup>+</sup> cells among total CD3<sup>+</sup> T cells.
